# Using a female-specific isoform of *doublesex* to explore male-specific hearing in mosquitoes

**DOI:** 10.1101/2025.03.09.642281

**Authors:** Matthew P Su, Marcos Georgiades, Marta Andrés, Jason Somers, Judit Bagi, YuMin M Loh, Yifeng YJ Xu, Kyros Kyrou, Andrea Crisanti, Joerg T Albert

**Affiliations:** Ear Institute, University College London, 332 Gray’s Inn Road, London, WC1X 8EE, UK; The Francis Crick Institute, 1 Midland Road, London, NW1 1AT, UK; Institute for Advanced Research, Nagoya University, Nagoya, 464-8601, Japan; Graduate School of Science, Nagoya University, Nagoya, 464-8601, Japan; Institute of Transformative Bio-Molecules (WPI-ITbM), Nagoya University, Nagoya, 464-8601, Japan; Animal Health Research Centre, National Institute for Agricultural and Food Research and Technology, Spanish National Research Council (CISA-INIA-CSIC), 28130 Valdeolmos, Spain; Department of Life Sciences, Imperial College London, UK; Department of Molecular Medicine, University of Padova, Padua, Italy; Cluster of Excellence Hearing4all, Sensory Physiology & Behaviour Group, Department for Neuroscience, School of Medicine and Health Sciences, Carl Von Ossietzky University Oldenburg, Carl Von Ossietzky Str. 9-11, 26111 Oldenburg, Germany

**Keywords:** *Anopheles gambiae*, *doublesex*, sex determination pathway, mosquito acoustic communication, hearing, ciliary function

## Abstract

Animal reproduction relies on elaborate divisions of labour and multiple dimorphisms between the sexes. Primary dimorphisms affect core elements of reproduction, secondary dimorphisms affect more indirect traits, including complex behaviours. In disease-transmitting mosquitoes, males locate females acoustically prior to copulation (phonotaxis). No comparable acoustic behaviour is known for females. As a result, the males’ ears – and hearing performance - have evolved to become substantially more complex. Sex-specific hearing in mosquitoes is in part controlled by the *doublesex* (*dsx*) gene. Intriguingly, *dsx* forms a linker between primary and secondary dimorphisms: spermatogenesis and ear morphogenesis share considerable molecular overlap and both depend on *dsx* expression patterns. We have combined transcriptomics with functional-anatomical analyses to dissect *dsx*-dependent hearing in the malaria mosquito *Anopheles gambiae*. By cross-linking our auditory findings to the genetic bases of spermatogenesis we advance the molecular understanding of sex-specific hearing mechanisms in insects, highlighting the special roles of ciliary factors therein.

**Highlights:**

- Disruption of the female-specific exon of the *doublesex* gene alters hearing in female *Anopheles gambiae* mosquitoes
- Female mutant ears are anatomically and functionally distinct from both wild-type male and female; they show a substantial, but incomplete, masculinisation
- Contrary to males, and similarly to females, the flagella of female mutants do not exhibit Self-Sustained Oscillations (SSOs)
- Transcriptomic comparisons of ears uncovered molecular bases of ‘auditory maleness’, these were closely linked to molecular machinery of ciliary function
- Comparisons with existing transcriptome from male testes revealed a substantial overlap between male ear and testicular ciliary motility machineries

## Introduction

In insects, sexually dimorphic behaviours are supported by sex-specific anatomical structures. An exemplary demonstration is the highly sexually dimorphic hearing system of male mosquitoes, whose function underlies mating. Anatomically, the male mosquito ear is a complex structure, containing many thousands of neurons and multiple populations of auditory efferents^1,2^. This male-specific complexity is required for successful copulation, as male mosquitoes detect and locate conspecific females within the crowded environment of (male-dominated) swarms^3^. Male phonotactic attraction to female flight sounds is augmented via spectral matching between female Wing Beat Frequencies (WBFs) and natural vibration frequencies of the male flagellar ear^4^. Phonotaxis appears highly conserved across multiple mosquito species^5^, though attempts at applying this knowledge for use in vector control have yet to be maximally exploited^6,7^.

In contrast to males, females from disease-transmitting mosquito species do not appear to demonstrate positive phonotaxis^8^. Immunological and neuroanatomical assays have enabled description of female auditory features, reporting a ∼50% reduction in the number of neurons and a more limited (or close-to non-existent in some species) efferent network as compared to males^1,9^. Despite this numerical reduction relative to males, the female mosquito ear still stands among the most complex insect hearing organs. Yet, female hearing-related behaviours have still to be elucidated, although numerous theories have been devised ranging from host seeking to predator avoidance to mate selection^5,8,10^. Likewise, the precise molecular substrates responsible for anatomical and behavioural differences between male and female ears remain unknown.

Beyond differences in auditory neuroanatomy and behaviour, there are clear dimorphisms in hearing function between male and female ears^9^. Male ears are tuned to higher frequency sounds than females at both mechanical and electrical levels^10^. Males also show a unique hearing state, referred to as Self-Sustained Oscillations (SSOs), in which the male flagellum – in the absence of any external stimulus - becomes an almost mono-frequent active oscillator with displacements orders of magnitude larger than in non-SSO states^9,11^. SSOs have been hypothesised to play a crucial role in mosquito courtship by entraining to, and enhancing, the flight tone of the female in the male ear^12^. Onset/offset of SSOs appears controlled by the efferent system and powered by active energy injection from JO neurons^9,11^. SSOs are restricted to males, with males from some species appearing to more readily demonstrate SSOs than others, though the anatomical and genetic factors which form their basis are unknown^9^.

Sex-specific alternative splicing of *doublesex* (*dsx*), the terminal molecular switch in the insect sex determination cascade, regulates somatic sexual differentiation (Figure 1A), giving rise to sexually dimorphic morphological features^13–15^. *dsx* plays an important role in sexual reproduction, and in particular the development of insect gonads^15–17^. Intriguingly, male sperm and mosquito auditory neurons are the only two specialised cell types that contain motile cilia, providing an interesting connection between two defining aspects of male mosquitoes (phonotaxis and sperm motility)^12,18^.

**Figure 1:**
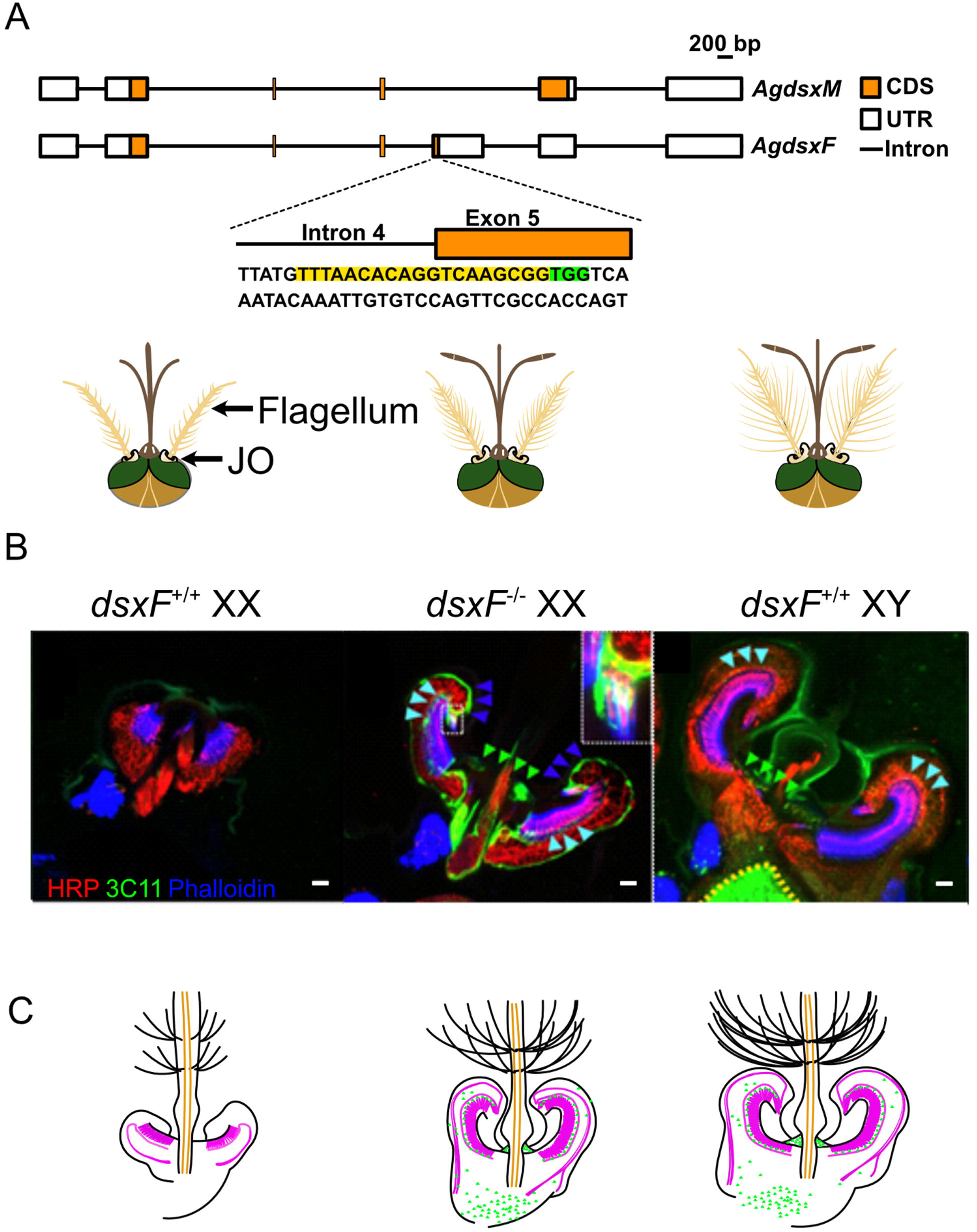
Altered auditory efferent innervation and JO size in *dsxF*^-/-^ female mutants. A) (*Above*) Schematic of male (M) and female (F) *Agdsx* isoforms (*AgdsxM* and *AgdsxF*, respectively); (*Below*) Schematic of *dsxF*^+/+^ *XX* (left), *dsxF*^-/-^ *XX* (centre) and *dsxF*^+/+^ *XY* (right) head anatomy based on previous publications. B) Horizontal sections of *dsxF*^+/+^ *XX* (left), *dsxF*^-/-^ *XX* (centre) and *dsxF*^+/+^ *XY* (right) JOs stained with presynaptic marker 3C11 (anti-synapsin, green) to label presynaptic efferent terminals within the JO and counterstained with neuronal marker anti-HRP (red) and phalloidin (blue) to label actin-based rods in scolopale and cap cells that surround auditory cilia. 3C11 also stains muscle motoneuron innervation in scape. Scale bar: 10 µm. Arrows highlight pre-synaptic efferent terminals picked up by 3C11 antibody. C) Schematic representations of each genotype’s JO anatomy based on B).

Previous work in *Anopheles gambiae* (*An. gambiae*) found that a loss-of-function mutation in the female-specific exon of *dsx* (hereafter referred to as *dsxF*) produces females with intersex morphological characteristics, including masculinisation of the external ear morphology and internal/external reproductive structures^15^. The shared motile cilia machinery between sperm and auditory neurons raises the question of how sex-specific isoforms of *dsx* differentially regulate the development of these tissues to support sex-specific reproductive and hearing behaviours. Whilst male ejaculate has been extensively researched to identify factors involved in regulating female behavioural changes post copulation^18,19^, the molecular factors of male hearing have remained largely unexplored. Identifying common genes shared between the two tissues could help find new effective vector control target genes that ideally simultaneously inhibit male hearing and reproduction.

This prior generation of an *An. gambiae dsxF* knockout mutant line offers an ideal tool for investigations of sexual dimorphisms in mosquito auditory systems^15^. In addition to the aforementioned intersex phenotypic differences reported for *dsxF*^-/-^ *XX* mutants^15^, the mutation produces an apparent dose-dependent masculinising effect on female WBFs^20^. Importantly, this *dsxF* mutant line has also been suggested for release as part of a gene-drive based mosquito control programme^15,21^. Given the importance of hearing for mosquito mating (and thus reproduction^22^) it is vital to test auditory function in these mutants prior to release, particularly in light of the aforementioned changes in WBF.

Here, we investigated the auditory anatomy and function in *dsxF*^-/-^ mutants to better understand the molecular mechanisms underlying sexual dimorphisms in mosquito hearing. We found significant differences in ear anatomy in *dsxF*^-/-^ *XX* mosquitoes as compared to both male and female controls, with mutants exhibiting an intersex phenotype in terms of efferent terminals in the JO, as well as pedicel size. Functional experiments found altered and intersex mechanical and electrical tuning frequencies in *dsxF*^-/-^ *XX* mutants, reflecting incomplete masculinisation. By performing RNA sequencing analyses of *dsxF*^+/+^ *XY*, *dsxF*^+/+^ *XX* and *dsxF*^-/-^ *XX* ears, we identified numerous promising gene candidates upregulated in *dsxF*^+/+^ *XY* and *dsxF*^-/-^ *XX* compared to *dsxF*^+/+^ mosquitoes which may underlie the auditory masculinisation. Notably, through intersectional analyses of differentially expressed genes, we identified factors that may influence SSOs.

Finally, our bioinformatic analyses showed that genes and Gene Ontological (GO) terms related to motile cilia machinery (microtubules, dyneins, etc.) appeared enriched in *dsxF*^+/+^ *XY* and *dsxF*^-/-^ *XX* ears as compared to *dsxF*^+/+^ *XX*. Given the importance of ciliary factors for sperm motility and the role of *dsx* in modulating both ear morphogenesis and spermatogenesis, we compared the auditory transcriptome with published testes data. Our analyses identified both tissue-specific ciliary factors and factors shared between the two tissues, suggesting that the basal molecular machinery for ciliary motility might be conserved between sperm and auditory neurons with tissue-specific adaptations being conferred by sets of specialized factors.

## Results

### dsxF^-/-^ XX mutant ears are significantly different in size and structure to both dsxF^+/+^ XY and dsxF^+/+^ XX ears

The mosquito ear is comprised of a feathery flagellum attached at the base to the Johnston’s organ (JO), the site of auditory mechanotransduction, which is housed in the pedicel (Figure 1A)^1,23,24^. The JO contains not only thousands of neurons but also sexually dimorphic efferent terminals originating from the brain^1^. Anatomical differences between the sexes underlie differences in function and thus behaviour. Previous reports have highlighted differences in external ear structure in *An. gambiae dsxF^-/-^ XX* mutants, including an increased pedicel size and density of fibrillar hairs^15,20^, but have not investigated potential changes in neuronal architecture.

To investigate changes in ear anatomy resulting from the *dsxF* mutation, we used immunohistochemistry to investigate JO neuroanatomy^2^. *dsxF*^+/+^ *XY* showed extensive efferent innervation in their JOs, whilst no such innervation was observable in *dsxF*^+/+^ *XX* JOs (Figure 1B). On the other hand, the anatomy of *dsxF*^-/-^ *XX* JOs was clearly distinct from female controls (Figure 1B). *dsxF*^-/-^ *XX* JOs contained not only many more neurons than in *dsxF*^+/+^ *XX* JOs, but also efferent terminals. Still, *dsxF*^-/-^ *XX* JOs remained distinct from control male JOs, with the number of neurons and the magnitude of the efferent populations reduced in *dsxF*^-/-^ *XX* mosquitoes (Figure 1B,C).

In addition to differences in JO anatomy and the density of fibrillar hairs, there was a significant difference between the female genotypes in terms of flagellar length (Figure S1A; Mann-Whitney test; p = 1.458x10^-5^). *dsxF*^+/+^ *XY* flagella were longer than both *dsxF*^+/+^ *XX* and *dsxF*^-/-^ *XX* flagella (Mann-Whitney tests; p < 2.2x10^-16^ for both comparisons).

### dsxF^-/-^ XX mutant ears have different mechanical tuning properties to dsxF^+/+^ XY and dsxF^+/+^ XX ears

Given the sizeable differences found in JO neuroanatomy, we next tested how these changes affected auditory function using Laser Doppler Vibrometry (Figure 2A). We investigated differences in frequency tuning and amplification in *dsxF*^-/-^ *XX* mosquito auditory systems as compared to *dsxF*^+/+^ *XX* and *dsxF*^+/+^ *XY* mosquitoes.

**Figure 2:**
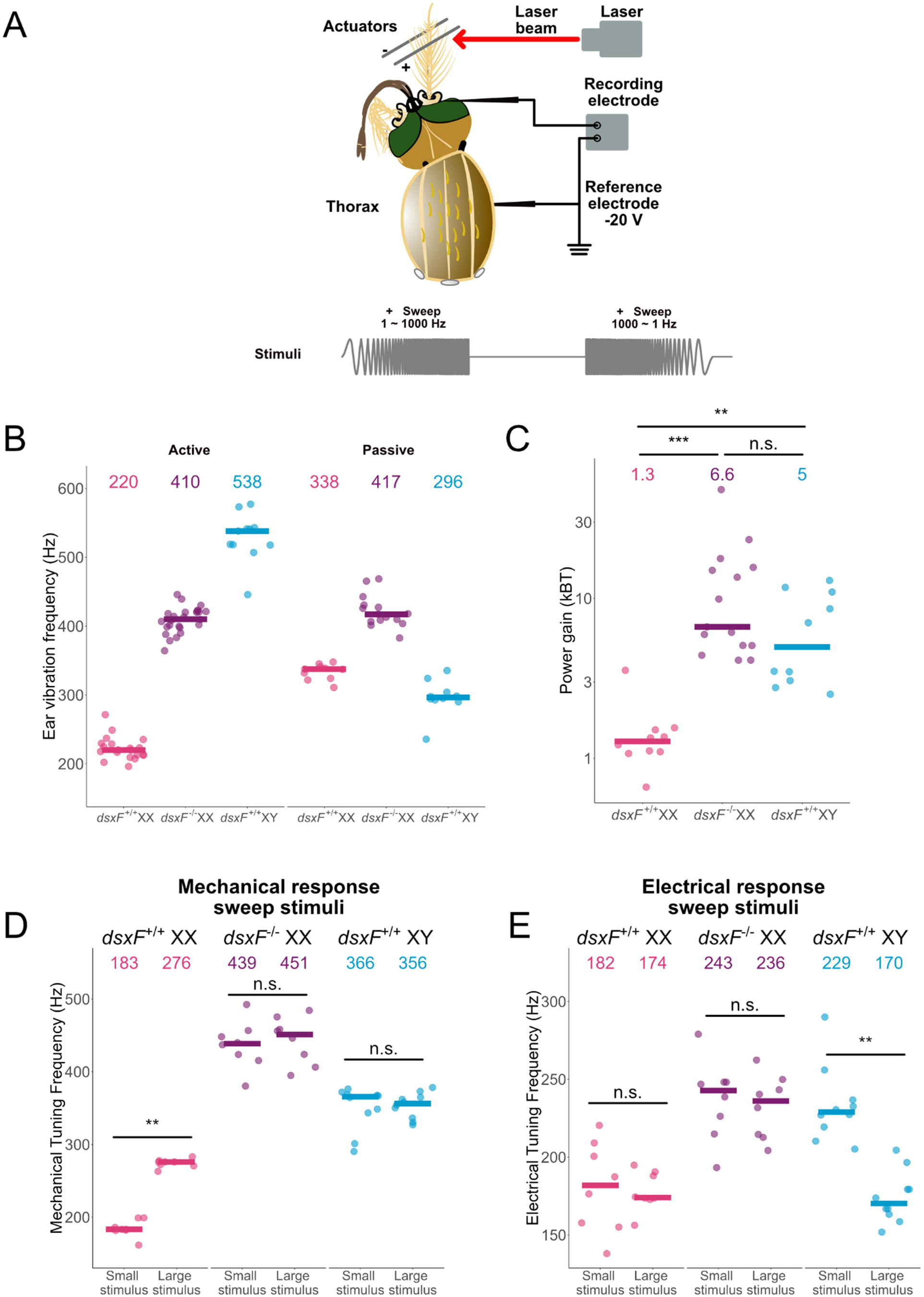
*dsxF*^-/-^ *XX* ears show differences in frequency tuning in both active and passive states compared to other *dsxF*^+/+^ *XX* and *dsxF*^+/+^ *XY* ears, as well as in electrical responses to stimulation. A) Experimental set-up for Laser Doppler Vibrometry (LDV) recordings. Laser beam is focussed towards the tip of the mosquito flagellum. Free fluctuation experiments used only unstimulated recordings. Experiments using electrostatic stimulation required the use of electrodes and electrostatic actuators. B) Calculated best mechanical tuning frequencies of mosquito flagellar receivers in active (left) and passive (right) conditions. Median values are represented by solid lines, which are printed for each group at the top of the panel. Individual points are best mechanical tuning frequencies for individual mosquitoes from each group. Sample sizes (females = active/ passive; males = active quiescent/ passive): *dsxF*^+/+^ □*XX* =□20/ 10; *dsxF*^-/-^□*XX* = 25/ 15; *dsxF*^+/+^□*XY* =□10/ 21. C) Calculated power gain for mosquito flagellar receivers. Median values are represented by solid lines, which are printed for each group at the top of the panel. Individual points are best mechanical tuning frequencies for individual mosquitoes from each group. All *dsxF*^+/+^□*XY* were in the non-SSO (quiescent) state for recordings. Mann-Whitney tests; ***, p<0.001; n.s., p>0.05. Sample sizes: *dsxF*^+/+^□*XX* =□10; *dsxF*^-/-^□*XX* = 15; *dsxF*^+/+^□*XY* =□10. D) Calculated best frequencies of mechanical responses to smallest and largest sweep stimuli for each genotype. Median values are represented by solid lines, which are printed for each group at the top of the panel. Individual points are best mechanical tuning frequencies for individual mosquitoes from each group. All *dsxF*^+/+^□*XY* were in the SSO state for recordings. Mann-Whitney tests; ***, p<0.001; n.s., p>0.05. Sample sizes: *dsxF*^+/+^□*XX* =□10; *dsxF*^-/-^□*XX* = 15; *dsxF*^+/+^□*XY* =□10. E) Calculated best frequencies of electrical responses to largest (top) and smallest (bottom) sweep stimuli for each genotype Median values are represented by solid lines, which are printed for each group at the top of the panel. Individual points are best mechanical tuning frequencies for individual mosquitoes from each group. All *dsxF*^+/+^□*XY* were in the SSO state for recordings. Mann-Whitney tests; ***, p<0.001; n.s., p>0.05. Sample sizes: *dsxF*^+/+^□*XX* =□10; *dsxF*^-/-^□*XX* = 15; *dsxF*^+/+^□*XY* =□10.

In the active state, *dsxF*^-/-^ *XX* mutants were found to be tuned to much higher frequencies than *dsxF*^+/+^ *XX* mosquitoes, with median frequencies of ∼410Hz as compared to 220Hz (Mann-Whitney test; p = 3.597x10^-8^; Table 1; Figure 2B). *dsxF*^-/-^ *XX* frequency tuning was also found to be significantly different from quiescent and SSO *dsxF*^+/+^ *XY* individuals (Mann-Whitney tests; p = 8.229x10^-6^ and p=1.386x10^-7^, respectively). Indeed, a frequency tuning of just over 400Hz places *dsxF*^-/-^ *XX* mutants between *dsxF*^+/+^ *XX* (∼220Hz) and non-SSO (quiescent) *dsxF*^+/+^ *XY* (∼520Hz) frequency tunings (Table 1).

**Table 1:**
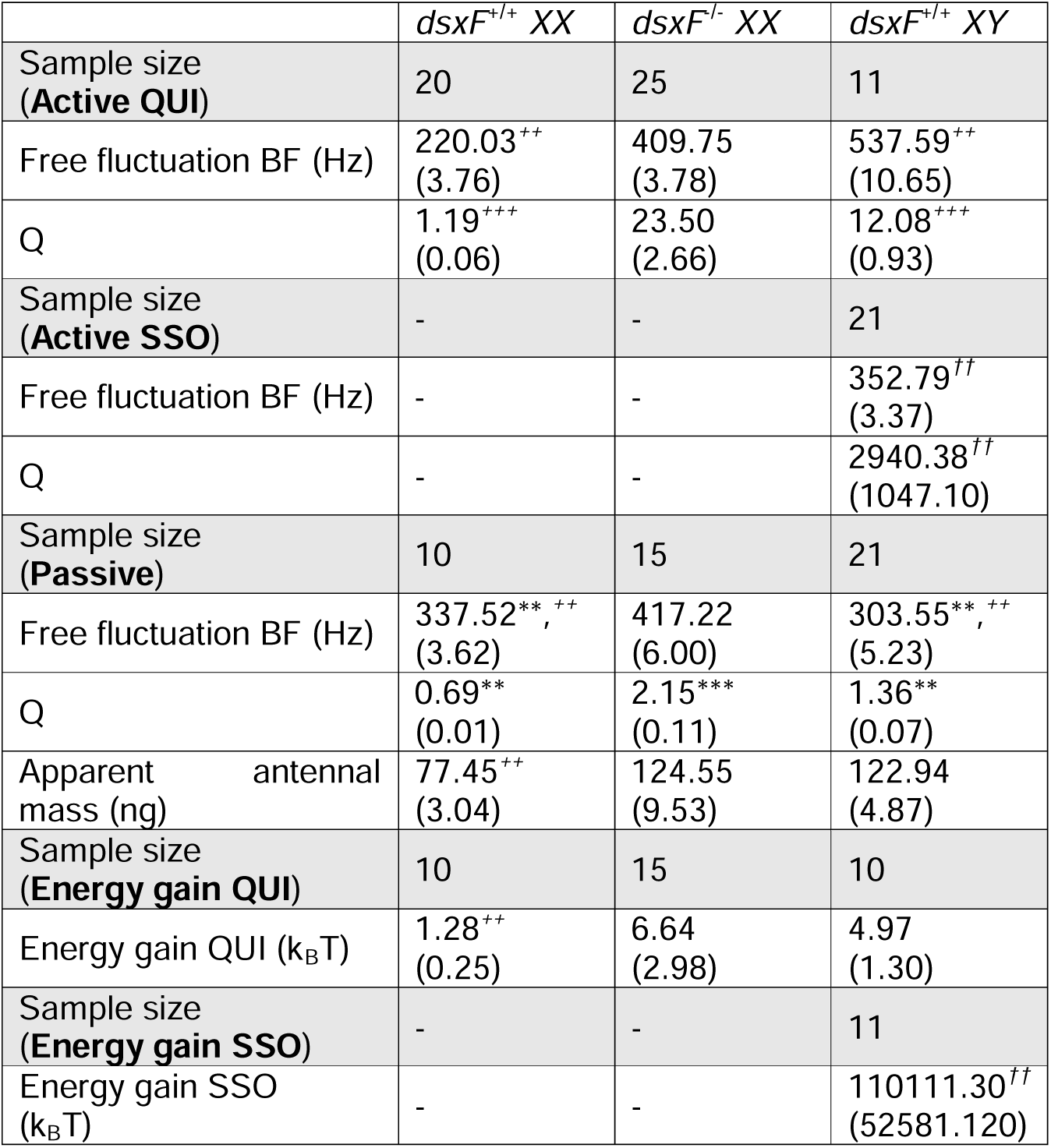
Parameters obtained from free fluctuation analyses for all groups. Median values from free fluctuation fits to data obtained from *dsxF^+/+^ XX* and *XY* mosquitoes, as well as *dsxF^-/-^ XX* mosquitoes, with SEM values provided in brackets. XX analyses include active and passive states, whilst XY active analyses divide males into either quiescent (QUI) or self-sustained oscillation (SSO) states. No significant differences between the fits for the passive state in QUI and SSO males were found for any parameter (Mann Whitney tests; p□>□0.05) so the results were combined. Parameters include the best frequency of the flagellum, tuning sharpness (Q), apparent antennal mass and estimated power gain for each group. Significant differences found between active and passive states within a specific mosquito group are starred (Paired Wilcox tests; **p□<□0.01, ***p□<□0.01). Significant differences between QUI and SSO dsxF^+/+^ XY in the active state are also highlighted (Mann Whitney tests; †p□<□0.05; ††p□<□0.01). Significant differences found in the best frequency, apparent antennal mass and energy gain between dsxF^-/-^ XX mosquitoes and all other mosquito groups are highlighted (Mann Whitney tests; +p□<□0.05; ++p□<□0.01; +++p□<□0.01).

| | $dsxF^{+/+}$ XX | $dsxF^{-/-}$ XX | $dsxF^{+/+}$ XY |
| --- | --- | --- | --- |
| Sample size<br>(Active QUI) | 20 | 25 | 11 |
| Free fluctuation BF (Hz) | 220.03<sup>++</sup><br>(3.76) | 409.75<br>(3.78) | 537.59<sup>++</sup><br>(10.65) |
| Q | 1.19<sup>+++</sup><br>(0.06) | 23.50<br>(2.66) | 12.08<sup>+++</sup><br>(0.93) |
| Sample size<br>(Active SSO) | - | - | 21 |
| Free fluctuation BF (Hz) | - | - | 352.79<sup>††</sup><br>(3.37) |
| Q | - | - | 2940.38<sup>††</sup><br>(1047.10) |
| Sample size<br>(Passive) | 10 | 15 | 21 |
| Free fluctuation BF (Hz) | 337.52<sup>**, ++</sup><br>(3.62) | 417.22<br>(6.00) | 303.55<sup>**, ++</sup><br>(5.23) |
| Q | 0.69<sup>**</sup><br>(0.01) | 2.15<sup>***</sup><br>(0.11) | 1.36<sup>**</sup><br>(0.07) |
| Apparent antennal mass (ng) | 77.45<sup>++</sup><br>(3.04) | 124.55<br>(9.53) | 122.94<br>(4.87) |
| Sample size<br>(Energy gain QUI) | 10 | 15 | 10 |
| Energy gain QUI ($k_B T$) | 1.28<sup>++</sup><br>(0.25) | 6.64<br>(2.98) | 4.97<br>(1.30) |
| Sample size<br>(Energy gain SSO) | - | - | 11 |
| Energy gain SSO ($k_B T$) | - | - | 110111.30<sup>††</sup><br>(52581.120) |

Notably, these differences extended into the passive state: whilst all other lines were found to have significant differences between active and passive states in terms of frequency tuning (Mann-Whitney tests; p = 3.579x10^-5^ and p= 0.0003669 for *dsxF*^+/+^ *XX* and *dsxF*^+/+^ *XY*, respectively), no such differences were found for *dsxF*^-/-^ *XX* mutants (Mann-Whitney test; p=0.4155).

*dsxF*^-/-^ *XX* mosquitoes also displayed significantly sharper frequency tuning, reflected in increased Q values, in the active state than any other lines (Mann-Whitney tests; p=3.609x10^-8^ and p= 0.0003591 for comparisons with *dsxF*^+/+^ *XX* and *dsxF*^+/+^ *XY*, respectively).

In terms of calculated energy gains in the quiescent state (QUIE), *dsxF*^-/-^ *XX* mutants displayed a masculine profile (Figure 2C), with energy gains similar to those exhibited by *dsxF*^+/+^ *XY*, and significantly greater to those exhibited by *dsxF*^+/+^ *XX* (Mann-Whitney tests; p=0.24 for *dsxF*^+/+^ *XY* compared to *dsxF^-/-^ XX*; p= 1.836x10^-6^ for *dsxF^-/-^ XX* compared to *dsxF*^+/+^ *XX*; p= 0.0006171 for *dsxF^+/+^ XY* compared to *dsxF*^+/+^ *XX*). Importantly, *dsxF*^+/+^ *XY* exhibited SSOs, a masculine trait never observed for either type of *XX* mosquito. SSOs were of lower frequencies than the mechanical tuning best frequencies of non-SSO (quiescent) *dsxF*^+/+^ *XY* individuals; the energy gain of SSO-ing *dsxF*^+/+^ *XY* flagellar ears was several thousand times greater than those calculated for any *dsxF*^-/-^ *XX* mutant mosquito (Fig S1B, C).

### dsxF^-/-^ XX mutant ears have different electrical tuning properties to dsxF^+/+^ XX and XY ears

We next used established electrostatic stimulation paradigms to combine our investigations into ear mechanical tuning with electrophysiological recordings focused on the frequency tuning of the auditory nerve^9^. By inserting an electrode into the antennal nerve, we were able to measure compound action potentials (CAPs) generated in response to electrostatic deflections of the mosquito flagellum. These deflections were induced by two stimuli types; force steps or pure tone sweeps.

First, we used step stimulation to investigate mechanical signatures of auditory transducer gating in both *XX* groups. *dsxF*^-/-^ *XX* mutants stiffness values were far greater than for *dsxF^+/+^ XX* individuals (Figure S2A; Table S1), and indeed greater than all previously reported mosquito stiffness values, including for males^9^. *dsxF*^-/-^ *XX* mutants also showed significantly greater CAP responses than *dsxF*^+/+^ *XX* for equivalent displacements (Figure S2B). Once these displacements were translated to the force domain, the significant extent of the increase in stiffness became apparent however, with greater forces required for *dsxF*^-/-^ *XX* mutants to elicit comparable CAPs to *dsxF*^+/+^ *XX* (Figure S2C).

The use of frequency sweeps, pure tones which linearly change in frequency from 1 to1000 or 1000 to 1Hz, facilitated investigation into peak mechanical and electrical tuning frequencies; that is, the stimulation frequencies at which the maximal flagellar vibrations and CAP responses were identified. Previous investigations found differences in female mechanical tuning frequencies across different stimulation amplitudes^9^; by changing the intensity of the stimulus we were able to test for frequency differences in both contexts across a range of stimulus amplitudes.

All *dsxF^+/+^ XY* mosquitoes exhibited SSOs throughout stimulus presentation. The mechanical best frequency of these SSO-ing *dsxF^+/+^ XY* mosquitoes did not change significantly when comparing the smallest to the largest stimulus intensities (paired Wilcox test; p=1; Table 2; Figure 2D), with best frequencies remaining between ∼345Hz and ∼365Hz across the entire range of stimulus intensities tested (Figure S2D). For *dsxF^+/+^ XX* mosquitoes however we saw an increase in best frequency from ∼190Hz to ∼275Hz from the smallest to the largest intensity, in agreement with previous reports^9^ of such a shift for white noise stimulation (paired Wilcox test; p = 0.007813; Figure 2D; Figure S2D). Similarly to *dsxF^+/+^ XY*, the mechanical best frequency of *dsxF^-/-^ XX* mutants did not change significantly when comparing the smallest to the largest stimulus intensities (439 Hz vs 451 Hz, respectively; paired Wilcox test; p=0.1953; Table 2; Figure 2D). *dsxF*^-/-^ *XX* mosquitoes maintained a mechanical best frequency near ∼445 Hz across the entire range of stimulus intensities (Figure S2D).

**Table 2:** Parameter values for each group for mechanical and electrical responses to sweep stimulation. Median values from analysis of mechanical and electrical responses to sweep stimulation *dsxF^+/+^ XX* and *XY* mosquitoes, as well as *dsxF ^-/-^ XX* mosquitoes, with SEM values provided in brackets.

| | $dsxF^{+/+}$ XX | $dsxF^{-/-}$ XX | $dsxF^{+/+}$ XY |
| --- | --- | --- | --- |
| **Sweeps** |  |  |  |
| Sample size | 8 | 8 | 10 |
| Mechanical BF largest sweep (Hz) | 276.04<br>(2.09) | 451.15<br>(11.33) | 356.26<br>(5.54) |
| Mechanical BF smallest sweep (Hz) | 183.45<br>(4.15) | 438.54<br>(11.53) | 365.75<br>(9.61) |
| Electrical BF largest sweep (Hz) | 174.03<br>(4.42) | 236.08<br>(7.18) | 170.25<br>(5.21) |
| Electrical BF smallest sweep (Hz) | 181.85<br>(10.23) | 242.78<br>(9.09) | 228.95<br>(7.71) |
| Displacement gain | 2.87<br>(0.15) | 1.38<br>(0.11) | 13.95<br>(0.97) |

On the other hand, the electrical best frequency of *dsxF*^+/+^ *XY* mosquitoes was found to decrease from ∼230Hz to ∼180Hz between the smallest and largest stimulus intensities, a phenomenon not observed for *dsxF*^+/+^ *XX* mosquitoes (paired Wilcox tests; p= 0.001953 and p= 0.7792, respectively; Table 2; Figure 2E). Similarly to females, the electrical best frequency of *dsxF*^-/-^ *XX* mutants did not change from the smallest to the largest intensity presentations (paired Wilcox tests; p=1; Table 2; Figure 2E). For these mutants, the electrical best frequency remained at ∼240Hz across the entire range of stimulus intensities tested (Figure S2D). In short, *dsxF*^+/+^ *XX* (female) nerves display only one uniform frequency tuning of ∼180Hz for both small and large stimulus intensities. *dsxF*^+/+^ *XY* (male) nerves, in turn, show two tuning regimes: for large stimulus intensities responses peak around ∼180Hz, resembling the tuning of female nerves but for small stimulus intensities an additional, higher-frequency tuning around ∼230Hz. The nerve responses of *dsxF*^-/-^ *XX* (intersex) females, finally, show the male-specific higher-frequency tuning at all stimulus intensities.

Changes in *dsxF*^+/+^ *XX* mechanical best frequency occurred in an almost step-like fashion once the stimulus force reached a specific threshold (Figure S2D). On the other hand, the shift in electrical best frequency observed for all male mosquitoes appeared more gradual, with a steady decrease in frequency for increasingly larger stimuli (Figure S2D).

We then calculated the displacement over force ratio for each stimulus intensity per individual mosquito and used these values to estimate the displacement gain for each genotype (Table 2). In contrast to the previous estimates of energy gain, we here found that *dsxF*^-/-^ *XX* mutants had significantly lower displacement gains than all other groups (Welch t-tests; p= 1.382x10^-6^ and p= 2.356x10^-5^ for comparisons with *dsxF*^+/+^ *XY* and *dsxF*^+/+^ *XX*, respectively). This finding is consistent with the vast increase in flagellar stiffness observed in intersex mosquitoes (Figure S2A; Table S1), which overshoots the male wildtype (*dsxF*^+/+^ *XY*) condition. Furthermore, *dsxF*^+/+^ *XY* mosquito displacement gain values were significantly greater than *dsxF*^+/+^ *XX* (Welch t-test; p= 3.339x10^-6^).

### RNAseq analyses of mosquito ears identified differences in expression of numerous ciliary motility related factor between groups

The above experiments focused on anatomical, and functional tests of hearing, demonstrating sizable differences between *dsxF*^-/-^ *XX* and control mosquitoes of both sexes, with several mutant characteristics shifting towards the *dsxF*^+/+^ *XY* mosquito phenotype. We investigated these differences at a transcriptional level to identify molecular factors that could account for the traits of ‘auditory maleness’.

Given previous reports of peaks in male and female *An. gambiae* activity at dusk^25^, we collected *dsxF*^+/+^ *XX* and *dsxF*^+/+^ *XY*, as well as and *dsxF*^-/-^ *XX*, mosquitoes at the onset of dusk, then dissected the pedicels of these mosquitoes and submitted them for RNA sequencing and analysis (Figure 3A). The following comparisons were conducted for differential gene expression:

**Figure 3:**
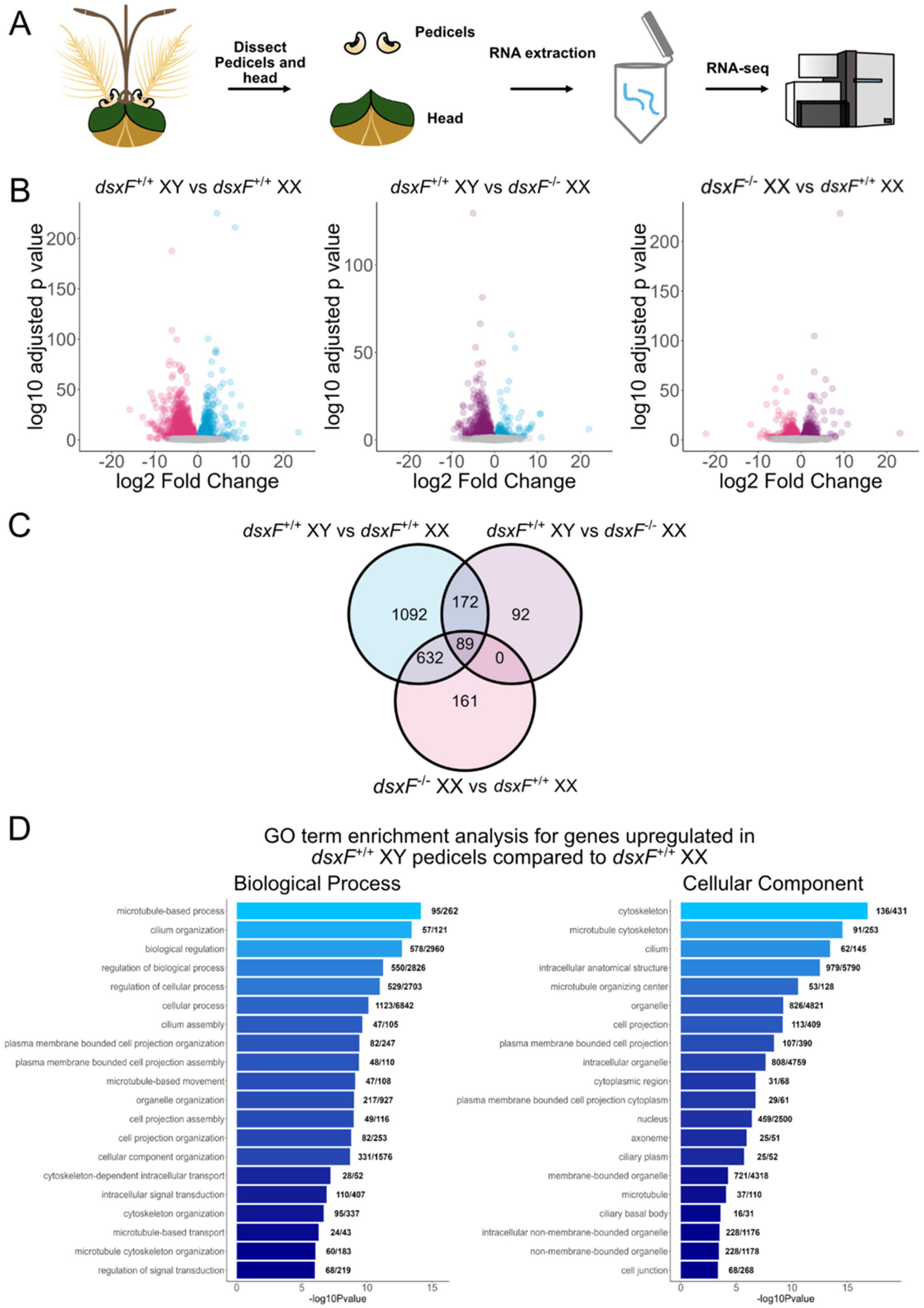
Significant enrichment of ciliary motility-related genes in *dsxF*^+/+^□*XY* **and *dsxF*^-/-^**□***XX* ears compared to *dsxF*^+/+^**□***XX***. A) Schematic of experimental assay, including tissue dissection. B) Volcano plots of differential gene expression between (left) *dsxF*^+/+^□*XY* and L*XX* pedicels, (middle) *dsxF* □*XY* and *dsxF* □*XX* pedicels and (right) *dsxF^-/-^*□*XX* and *dsxF*^+/+^□*XX* pedicels. C) Venn Diagram of pedicel gene intersections showing those differentially regulated between groups. Upregulated group is to left hand side of group name. D) GO term enrichment analysis for genes upregulated in *dsxF*^+/+^□*XY* pedicels compared to *dsxF*^+/+^□*XX* (from Table 3). Left, Biological Processes; Right, Cellular Component.

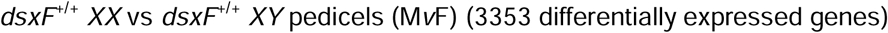

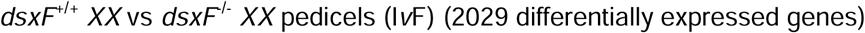

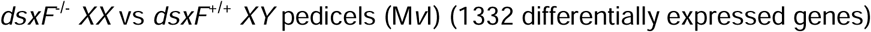

These comparisons are visualized *via* Volcano plots (Figure 3B) and a Venn diagram (Figure 3C). Raw count data is provided in Supplemental File 1. To explore the molecular underpinnings of auditory maleness, from the 3353 differentially expressed genes in the *dsxF*^+/+^ *XY* vs *dsxF*^+/+^ *XX* (denoted M*v*F) comparison, we focused our primary analyses on the 1985 genes found to be significantly upregulated in male pedicels as compared to females. We used the *dsxF*^-/-^ *XX* vs *dsxF*^+/+^ *XX* (denoted I*v*F) and *dsxF*^+/+^ *XY* vs *dsxF*^-/-^ *XX* (denoted M*v*I) comparisons to create subsets of the 1985 genes upregulated in the M*v*F comparison. Of the 1985, 632 genes (subset *s_632_*) were found to be upregulated in both *dsxF*^+/+^ *XY* and *dsxF*^-/-^ *XX* compared to *dsxF*^+/+^ *XX*, but were not differentially expressed between *dsxF*^+/+^ *XY* and *dsxF*^-/-^ *XX*. Thus, the subset contains genes highly expressed in *dsxF*^+/+^ *XY* compared to *dsxF*^+/+^ *XX* that also exhibit full recovery of male expression in *dsxF*^-/-^ *XX*.

One hundred and seventy two genes (subset *s_172_*) were found to be upregulated in *dsxF*^+/+^ *XY* compared to *dsxF*^+/+^ *XX* and *dsxF*^-/-^ *XX*, but not differentially expressed between the *dsxF*^-/-^ *XX* and *dsxF*^+/+^ *XX* genotypes. These genes thus exhibit higher expression in *dsxF*^+/+^ *XY* and are not under the influence of the DsxF mutation. Of the remaining genes, 89 (subset *s_89_*) were found to be upregulated in *dsxF*^+/+^ *XY* compared to both *dsxF*^+/+^ *XX* and *dsxF*^-/-^ *XX*, but were also upregulated in *dsxF*^-/-^ *XX* compared to *dsxF*^+/+^ *XX*. Thus, this subset represents transcripts highly expressed in *dsxF*^+/+^ *XY* in which *dsxF*^+/+^ *XY* expression has been recovered only partly in the *dsxF*^-/-^ *XX* group.

Finally, the last 1092 genes (subset *s_1092_*) of the 1985 set were found to be upregulated in *dsxF*^+/+^ *XY* compared to *dsxF*^+/+^ *XX* but not in *dsxF*^+/+^ *XY* compared to *dsxF*^-/-^ *XX* nor in *dsxF*^-/-^ *XX* compared to *dsxF*^+/+^ *XX*. This large subset includes genes that, on average, showed increased expression in *dsxF^-/-^ XX* compared to *dsxF^+/+^ XX* but not enough to yield statistically significant results. Subsets of the 1985 genes are shown in Figure 3C; Figure S3 includes visual comparisons of the log median expression for each genotype within each subset to aid understanding of their interpretations. Full gene lists for each subgroup are included in Supplemental File 2.

These subsets thus facilitate informed explorations into DsxF- and non-DsxF-inhibited auditory maleness, and also inferences on the molecular basis of our experimental findings. The transcripts of *s_632_* constitute the transcriptomic basis of auditory maleness under the inhibition of DsxF, whilst those of *s_89_* reflect those aspects of auditory maleness only partly influenced by it; transcripts of *s_1092_* may also reflect these components at a level non-differentiable by statistical analysis. Finally, the transcripts of *s_172_* constitute auditory maleness that is not under the regulation of DsxF.

We next conducted global, gene ontology (GO) enrichment, and gene-specific annotation analyses independently on each of the three sets to investigate these spaces. Selected elements of GO enrichment analyses are included in Supplemental File 3 and their expression levels visualised in Figure 3D.

GO analyses on *s_632_* revealed several terms relating to the structure and organisation of microtubules, cilia, and the motor complex. These include microtubular structural components such as AGAP008275-RA (tektin-2), AGAP000219-RA (tektin-3), AGAP001227-RA (gamma-tubulin complex component 5), and AGAP010971-RA (tubulin), AGAP002334-RA (spastin), which functions as a microtubule severing protein, AGAP006324-RA (*nompB*), which is required for the assembly of sensory cilia, AGAP005250-RA (protein kintoun) and AGAP009594-RA, which are dynein assembly factors. Other proteins related to microtubule and ciliary structure and regulation highlighted by our analyses are AGAP012726-RA and AGAP003120-RA, which are stabilisers of axonemal microtubules, AGAP013430-RA (piercer of microtubule wall), and AGAP006735-RA and AGAP008870-RA (cilia and flagella associated proteins). Thus, at least partly, DsxF-inhibited auditory maleness seems to involve setting up the structures used by motor proteins to generate motion. In terms of transcripts encoding motor proteins, *s_632_* included some kinesins (AGAP007592-RA, AGAP002427-RA, AGAP000561-RA) and dyneins (AGAP011441-RA, AGAP000320-RA, AGAP010165-RA, AGAP009568-RA, AGAP004416-RA).

We saw previously that both *dsxF^+/+^ XY* and *dsxF^-/-^ XX* mosquitoes exhibited similar power gains that were also greater than *dsxF^+/+^ XX* mosquito estimates; the shared structural and motor elements just discussed may directly relate to this finding. We also saw that contrary to *dsxF^+/+^ XX* mosquitoes, both *dsxF^+/+^ XY* and *dsxF^-/-^ XX* mosquitoes have mechanical frequency tuning largely invariant to stimulus intensity. Set *s_632_* includes elements related to ion channels, receptors and signalling that could be involved in tuning the flagellum. The set includes, for example, *nanchung* (AGAP012241-RA), a key element of the mechanotransduction complex. Further elements of *s_632_* include *AgOct*β*2R* (AGAP002886-RA), an octopamine receptor that has recently been shown to influence male ear tuning^12^, a piezo-type mechanosensitive ion channel (AGAP009276-RB), some chloride ion channels (AGAP005599.R500, AGAP005599.R502, and AGAP009616-RA), some potassium ion channels (AGAP001281-RB, AGAP003709-RC, AGAP003709-RD), a calcium ion channel (AGAP008028-RA) and a G protein-coupled receptor (AGAP001562-RA). Finally, several cuticle-related transcripts (AGAP000986-RA, AGAP000988-RA, AGAP006830- RA, AGAP006828-RA, AGAP009868-RA, AGAP006283-RB, and AGAP012487-RA) were included in *s_632_* and could be involved in the male-like enlargement of the JO observed in *dsxF^-/-^ XX* mutants.

As alluded to above, transcripts related to auditory maleness only partly influenced by DsxF are included in *s_89_*, and possibly also *s_1092_*. Several ion channels, receptors and signalling related transcripts that were only partly recovered in the *dsxF^-/-^ XX* mutants relative to *dsxF^+/+^ XY* are found in *s_1092_*, including two potassium ion channels (AGAP000254-RA and AGAP011924-RA), the GABA-gated chloride ion channel *Rdl* (AGAP006028-RC) and three other chloride ion channels (AGAP005599.R499, AGAP005599.R504, and AGAP005777-RA), *TRPM* (AGAP006825-RA), a nicotinic acetylcholine receptor component (AGAP002152-RA), two ionotropic receptors (AGAP001478-RA and AGAP005527-RA), and two G protein-coupled receptors (AGAP002888-RC and AGAP004222-RA).

Set *s_89_* includes just two transcripts encoding dynein motor proteins (AGAP008689-RB and AGAP011540-RA) that were only partly recovered in *dsxF^-/-^ XX* mutants relative to *dsxF^+/+^ XY*, while set *s_1092_*includes two dyneins (AGAP006887-RA and AGAP004030-RA), two kinesins (AGAP000159-RA and AGAP000575-RA), and four myosins (AGAP004403-RA, AGAP000776-RA, AGAP029989.R839, and AGAP011138-RB). A characteristic male feature that was not recovered in *dsxF^-/-^ XX* was that of self-sustained oscillations (SSOs) of the flagellum. It is possible that partial recovery of expression in the signalling and motor-related transcripts just discussed is related to the lack of SSOs in the *dsxF^-/-^ XX* mutants. These transcripts, thus, may form part of the machinery that brings SSOs about.

It is also possible that SSOs are a manifestation of auditory maleness that is not under the influence of DsxF. Looking at *s_172_* we find several ion channels, receptors and signalling transcripts that are significantly higher in *dsxF^+/+^ XY* compared to both *dsxF^+/+^ XX* and *dsxF^-/-^ XX* but similar in expression in *dsxF^+/+^ XX* and *dsxF^-/-^ XX*. Such transcripts could be key factors for SSOs, as these have been shown to be influenced by neurochemical and efferent-related modulation of the male hearing system^9,12^. The set includes two potassium ion channels (AGAP001284-RA, AGAP005251-RA), a histamine-gated chloride ion channel subunit (AGAP001990-RA), a nicotinic acetylcholine receptor subunit (AGAP005034-RA) and a metabotropic glutamate receptor (AGAP005034-RA).

### Comparison with previously published testes RNAseq data highlights tissue-specific factors related to the motile cilia machinery

The presence of SSOs in male mosquitoes constitutes complementary evidence for their “active hearing”, the broader phenomenon by which the ear injects energy into and thereby amplifies the reception of external signals. Several motor protein and motor protein-related transcripts were identified in the previous paragraphs that characterise both DsxF-inhibited and DsxF-independent male-specific expression, suggesting a more fundamental link between auditory cellular motility and maleness as auditory neurons and sperm are the only two known specialised cells that contain motile cilia in insects (Figure 4A).

**Figure 4:**
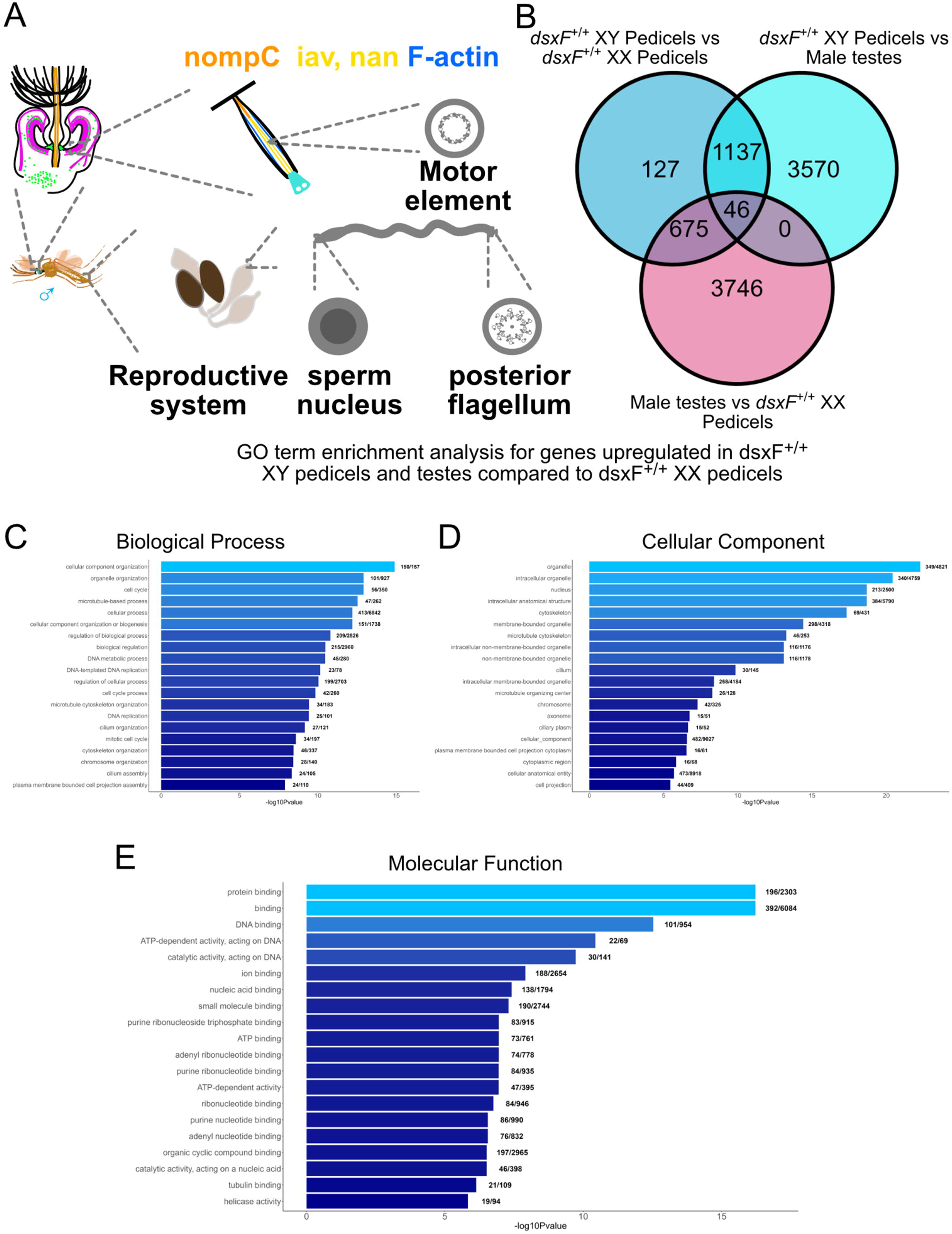
Comparisons of ears and testes enables identification of tissue specific ciliary motility-related factors. A) Schematic of tissues containing motile cilia in male mosquitoes. B) Venn diagram of pedicel/testes gene intersections showing those differentially regulated between groups. Upregulated group is to left hand side of group name. C) GO term enrichment analysis (Biological Process terms) for genes upregulated in male pedicels and testes compared to *dsxF*^+/+^□*XX*. Upregulated group is to left hand side of group name. D) GO term enrichment analysis (Cellular Component terms) for genes upregulated in male pedicels and testes compared to *dsxF*^+/+^□*XX*. Upregulated group is to left hand side of group name. E) GO term enrichment analysis (Molecular Function terms) for genes upregulated in male pedicels and testes compared to *dsxF*^+/+^□*XX*. Upregulated group is to left hand side of group name.

To investigate this relationship further we next contrasted our pedicel RNAseq dataset with an existing testes RNAseq database^26^. We identified 675 genes (*s_675_*) upregulated in both male pedicels and testes compared to *dsxF*^+/+^ *XX* pedicels (Figure 4B). GO enrichment analyses applied to *s_675_* genes identified terms related to dynein assembly and microtubule binding/regulation (Figure 4C, D and E). These dynein and microtubule-related genes could also be differentially regulated between male tissues, with some showing significantly increased expression only in pedicels or testes (Supplemental File 4). Further analyses showed that, similar to *s_632_* of the pedicel comparisons intersectional subsets, *s_675_* included transcripts AGAP005250-RA (protein kintoun), AGAP010971-RA (tubulin) and AGAP008275-RA (tektin-2).

Further, *s_675_* included two ionotropic receptors also found in *s_1092_* (AGAP005527-RA and AGAP001478-RA), a potassium ion channel (AGAP003709-RA) and a piezo-type mechanosensitive ion channel component (AGAP009276-RB) that were also in *s_632_*. In terms of motor-related transcripts *s_675_* included one dynein (AGAP004030-RA), two myosins (AGAP004403-RA and AGAP01138-RA) and one kinesin (AGAP000575-RA) that were also in *s_1092_* (with the exception of AGAP01138-RA; *s_1092_* included the - RB isoform of the transcript). Set *s_675_* also included four additional dyneins that were in *s_632_* (AGAP004416-RA, AGAP011441-RA, AGAP009568-RA, and AGAP010165-RA).

## Discussion

### Disruption of the female specific exon of doublesex leads to female hearing systems being partially converted to male hearing systems

Sexual dimorphisms in mosquito hearing function are in part the result of differences in ear anatomy. The roles played by auditory efferents are a major part of these differences in function^1^. Auditory efferents, though clearly abundant in the ears of male *An. gambiae*, were found in far smaller numbers for conspecific females both here and in previous work^9^. The *dsxF* loss-of-function mutation appeared to not only increase the number of neurons in mutant female ears compared to wild-type females, but also to increase the number of efferents (Figure 1). This mutation was not sufficient however to completely transform female ears into male ears, with *dsxF*^-/-^ *XX* ears remaining smaller than all male ears in terms of neuronal counts.

Functionally, *dsxF*^-/-^ *XX* mutant mechanical and electrical tuning appeared different from controls of both sexes (Figure 2, Figure S2). Mechanical frequency tuning was distinct from both control males and females (Table 1). *dsxF*^-/-^ *XX* mutants were able to produce significantly greater compound action potentials than *dsxF*^+/+^ *XX* following exposure to identical stimuli in terms of flagellar displacement, in agreement with their apparent increase in neuronal number. Interestingly, the *dsxF*^-/-^ *XX* ear was also found to be much stiffer than ears of *dsxF*^+/+^ *XX*, with substantially greater amounts of force required to achieve equivalent displacements (Figure S2). Neurons have been shown to contribute to the stiffness of the antennal sound receiver in *Drosophila*^27^; the increase of neuronal number in *dsxF*^-/-^ *XX* ears is thus a contributing cause for the increased CAP responses and stiffness, but other causal factors are also likely to be at play. Increases in stiffness could have several underlying causes, from changes in ion channel expression^28^ to altered cuticular structures attaching JO neurons to the flagellum^29,30^ (Supplemental File 2). Most notably in this context, another principal parameter of receiver mechanics, i.e. the apparent mass of the *dsxF*^-/-^ *XX* flagellum is fully restored to the level of the *dsxF*^+/+^ *XY* controls.

### SSOs remain a male-specific phenomenon

Perhaps most importantly, however, mutants failed to exhibit SSOs, the hallmark of male mosquito ears and potentially the central component of the male’s ability to identify the faint sound of a female within the din of a swarm^9,11,12^. Autonomous receiver vibrations have been observed in females following injection of DMSO^11^, but these are quantitatively and qualitatively distinct from the SSOs seen in male receivers, and much more resemble the autonomous oscillations observed in *Drosophila* after genetic or pharmacological manipulation. Most notably, injection of compounds, which specifically severe the efferent/afferent feedback loops between the ear and brain induced SSOs only in males^9^. Our data – and our corresponding intersectional analysis - will help to close in on the molecular machinery for SSOs. While it is possible, and even likely, that SSOs originate from the interaction of several molecular factors, our study has made a dedicated effort to narrow the list of candidates down to a manageable number. The most promising list is provided by subset *s_172_*.

The substantial amplification (lifting the receiver’s power gain by almost 4 orders of magnitude) provided by the male’s SSOs is a remarkable feature^9^, rivalling the unique amplification carried out by the piezo-crystal-like protein Prestin in the lateral walls of mammalian outer hair cells^31^. While it is likely that SSOs originate from ciliary motility factors, such as dyneins, and while it has also been shown that the auditory power gain in *Drosophila* is independent of Prestin homologues^32^, it remains to be seen if this also holds true for mosquitoes. Although the precise molecular causes of SSOs are thus not yet resolved, it seems clear that SSOs are a property that originates from the auditory JO neurons proper, though they may be modulated by efferent, neuromuscular or other synaptic input^33^.

### RNAsequencing analyses identified changes in multiple motile ciliary related factors

Our RNAseq analyses found upregulation of a number of ion channels and receptors in *dsxF*^+/+^ *XY* pedicels as compared to *dsxF*^+/+^ *XX*. Any one, or combination, of these candidates may be necessary for SSOs (Table 3). Beyond changes in ion channel counts however, our analyses also found substantial enrichment of motile ciliary-related factors in *dsxF*^+/+^ *XY* compared to *dsxF*^+/+^ *XX* pedicels some of which were fully recovered by the DsxF mutation and some only partly – this extended to microtubules, dyneins, myosins, kinesins and flagella-associated proteins.

These fundamental changes in the motile apparatus underlying active hearing function may offer new avenues to explore sexual dimorphisms in the fundamental ear machinery. Our findings highlight the developmental and mechanistic relationship between male ears and testes, with several motor proteins, and other cell motility related elements being co-expressed in the two tissues (Supplemental File 4). In particular, several motility-related transcripts that reached “masculine” (or towards masculine) levels of expression in the ears of *dsxF^-/-^ XX* mutants, showed shared levels expression between male ears and testes and had significantly higher expression than the respective transcripts in the female ear. This finding is also promising in terms of vector control interventions targeting reproduction as disruption of molecular elements expressed in both tissues could improve the potency of interventions by disrupting both hearing function and sperm motility^34,35^.

### Investigating the sex determination pathway facilitates identification of novel targets for vector control

*dsx^35,36,37^* represents only one component of the mosquito sex determination pathway, with *femaleless, SOA* and *yob* all playing separate roles^13,36–39^. Considering that many of these lines have been highlighted for use in mosquito control programmes, it is essential that their auditory function be first tested before release not only in *Anopheles* but also for other species. The mosquito ear represents a highly promising target for novel vector control methods, with the development of insecticidal resistance increasing the pressure to develop new control options^5,40^. The overwhelming reliance of males on hearing for the identification of conspecific females during courtship suggests that interfering with this process via pharmacological or acoustic tools holds great potential. However, acoustic lures targeting males thus far have failed to transfer their efficacy from the lab to field trials^41^; identifying the crucial component of this failure is challenging without first understanding the networks which underlie mosquito hearing.

### Limitations of our study

Here, we investigated only the effect of disrupting the female specific isoform of *dsx* (*dsxF*); the role played in the sex determination pathway by the male specific isoform (*dsxM*) therefore remains unclear. We were unable to identify the precise molecular basis of male mosquito SSOs, though our gene lists provide guidance for future research. Finally, whilst *dsx* plays major roles during development, in this study we investigated only adult mosquito ear anatomy and gene expression; future work could explore the effect of DsxF mutation on the development of *An. gambiae* JOs.

## Acknowledgements

The authors would like to thank Carla Siniscalchi (Imperial College London) for providing G3 strain pupae.

This work was supported by a grant from the Biotechnology and Biological Sciences Research Council, UK (BBSRC, BB/V007866/1 to J.T.A.), a grant from The Human Frontier Science Program (HFSP grant RGP0033/2021 to J.T.A.) and from the European Union’s Horizon 2020 research and innovation programme (H2020-ERC-2014-CoG/648709/Clock Mechanics, to J.T.A.). M.P.S. and J.S. were supported by the European Research Council grant H2020-ERC-2014-CoG/648709, an ANTI-VeC pump-priming grant (AV/PP/0028/1, to J.T.A.) and a UCL GCRF small grant (QR GCRF 551039 to J.T.A.). M.G. was also supported by the UCL GCRF small grant and the ANTI-VeC pump-priming grant. M.P.S, Y.M.L. and Y.Y.J.X. were co-funded by a Tokai Pathways to Global Excellence (T-GEx), as part of the MEXT Strategic Professional Development Program for Young Researchers, grant (0121an0002), and an MEXT KAKENHI Grant-in-Aid for Research Activity Start-up (No. JP22K15159). M.A. received funding through a Marie Sklodowska-Curie Individual Fellowship from the European Commission (H2020-MSCA-IF-2016/752472). J.B. was supported by BB/V007866/1. J.T.A. was supported by the European Research Council grant H2020-ERC-2014-CoG/648709, UCL GCRF and ANTI-VeC grants. RNAseq was provided via InfraVec2 (#5251).

## Contributions

Conceptualization, M.P.S., M.G., M.A., J.S., K.K., A.C. and J.T.A.; Methodology, M.P.S., M.G., Y.M.L., Y.Y.J.X. and J.T.A.; Investigation, M.P.S., M.G., M.A. and J.B.; Resources, M.P.S. and J.T.A.; Writing, M.P.S., M.G., Y.M.L. and J.T.A.; Funding Acquisition, M.P.S. and J.T.A.; Supervision, M.P.S. and J.T.A.

## STAR Methods

### Key resources table

**STAR Methods:** **Key resources table**
| REAGENT or RESOURCE | SOURCE | IDENTIFIER |
| --- | --- | --- |
| **Antibodies** |  |  |
| Mouse monoclonal anti-SYNORF1 3C11 | Developmental Studies Hybridoma Bank | AB_528479 |
| Cy<sup>TM</sup>3 AffiniPure<sup>TM</sup> Goat Anti-Horseradish Peroxidase | Jackson Immuno Research | AB_2338959 |
| Phalloidin, Alexa Fluor<sup>TM</sup> 647 | Thermo Fisher | #A-22287 |
| Goat anti-mouse IgG Secondary Antibody, Alexa Fluor<sup>TM</sup> 488 | Thermo Fisher | #A-11029 |
| **Deposited data** |  |  |
| RNAsequencing data | This paper | GEO: GSE289862 |
| **Experimental models: Organisms/strains** |  |  |
| *Anopheles gambiae* (G3 strain) | Kyrou et al<sup>15</sup> | N/A |
| *Anopheles gambiae* (doublesex<sup>F</sup> mutant strain) | Kyrou et al<sup>15</sup> | N/A |
| **Software and algorithms** |  |  |
| ImageJ (version 1.54g) | ImageJ | https://imagej.net/ij/ |
| VibSoft (version 6.0) | Polytec | https://www.polytec.com |
| R (version 4.3.1) | R Core Team | https://www.R-project.org/ |
| Spike2 (version 10.08) | Cambridge Electronic Design | https://ced.co.uk/products/spkovin |

## Resource availability

### Lead contact

Further information on all experiments conducted as part of this report, in addition to requests for resources, can be requested from the lead contact, Joerg T. Albert.

### Materials availability

This study did not generate new unique reagents.

### Data and code availability

- RNAsequencing data is available via the lead contact upon request.
- This paper does not report original code.
- Additional information regarding analysis protocols/data collection is available via from the lead contact upon request.

### Experimental model

*Mosquito rearing*: *An. gambiae* G3 strain (*dsxF*^+/+^) and *dsxF*^-/-^ *XX* mutant pupae were reared by the Crisanti lab at Imperial College London.

*dsxF*^+/+^ pupae were sex separated and kept in single sex cages in incubators maintained at 27°C and 60-70% relative humidity using a 12 hour:12 hour light/ dark cycle. *dsxF*^-/-^ pupae were not sex separated but were otherwise reared in identical conditions. Mosquitoes were supplied with a constant source of 10% glucose solution. Cow blood feeding, when required, was conducted by a trained research assistant using a Hemotek system (Discovery Workshops, Accrington).

All mosquitoes used for experiments, unless otherwise noted, were virgin and aged 3 – 6 days old.

### Method details

*Immunohistochemistry*: Samples were prepared following previously published protocols^9,42^. Heads were removed from adults from each genotype and fixed in 4% paraformaldehyde for anLhour at room temperature; their proboscises were also removed prior to fixation. Following this, heads were embedded in albumin/gelatin and post-fixed overnight in 6% formaldehyde at 4□°C. Each block was then sectioned using a vibrotome into 40□µm sections and washed in phosphate-buffered saline 0.3% Triton X-100. Samples were blocked in 5% normal goat serum and 2% bovine serum albumin.

Primary antibodies used included: monoclonal antibody 3C11 (anti-SYNORF1; 1:50; Developmental Studies Hybridoma Bank, University of Iowa, http://dshb.biology.uiowa.edu/) and conjugated primary antibody anti-HRP-Cy3 (1:500, Jackson ImmunoResearch, Code: 123-165-021). Secondary antibodies used included corresponding Alexa Fluor Dyes (1:500; Thermo Fisher). Sections were mounted onto slides using DABCO and then visualised using a Zeiss 880 confocal microscope.

*Flagellar length measurements*: The right flagellae of adult mosquitoes from each genotype were removed using a pair of forceps whilst the mosquitoes were CO_2_ sedated. The flagellae were then transferred to separate microscope slides in groups of five. Each individual sample was then immediately imaged using a Zeiss Axioplan 2 microscope and Axiovision 4.3 software. Wing lengths were determined using Axiovision 4.3 software length measurement function, calibrated to the nearest 0.1 mm. Three biological repeats were conducted over different generations.

Total sample sizes for each group: *dsxF*^+/+^ *XX* = 59; *dsxF*^-/-^ *XX* = 40; *dsxF*^+/+^ *XY* = 56.

*Laser Doppler Vibrometry - Mosquito preparation*: As described previously, mosquitoes were glued to a Teflon rod using blue-light-cured glue^9^. Glue was applied across the mosquito body to restrict movement, and thus disturbances, during recordings. The left flagellum was glued to the head before glue was applied between the pedicels. Only the right flagellum therefore was able to move completely unhindered.

The mounted mosquito was held in place by a micromanipulator placed on a vibration isolation table opposite a PSV-400 laser Doppler vibrometer (Polytec) with an OFV-70 close up unit and a DD-500 displacement decoder. The mosquito was orientated so as to face the laser Doppler vibrometer at a 90° angle. The laser was focussed on the second flagellomere from the flagellum tip for all males, whilst the third flagellomere from the tip was used for all females. All LDV experiments took place in a temperature-controlled room (21-23°C).

*Laser Doppler Vibrometry - Force step and sweeps procedure*: A reference electrode was inserted into the thorax of a prepared mosquito to raise its’ electrostatic potential to −20 V relative to ground. A pair of electrostatic actuators were then positioned symmetrically around the unrestricted flagellum to enable electrostatic stimulation of the ear. A recording electrode was inserted at the base of the right pedicel to allow for recording of compound antennal nerve responses to the electrostatic stimulation.

The actuators were then used to provide stimulation to the flagellum. Flagellar displacement was monitored using the LDV, with simultaneous electrophysiological activity also recorded. Free fluctuations of the flagellum were recorded at the beginning and end of all experiments to assess the status of the mosquitoes’ auditory system.

Force step stimuli provided were identical to those reported previously^9^ and increased in intensity in logarithmic steps; 20 different intensities were used for each experiment (i.e. 20 different step magnitudes). Sweeps consisted of a pure tone stimulus which either increased linearly from 1 to 1000Hz over the course of 1 second (denoted as ‘forward sweep’), or decreased linearly from 1000 to 1Hz over the same time frame (denoted as ‘backward sweep’). Forward and backward sweeps were played sequentially, with a 1 second, unstimulated, pause in between each sweep. 10 different sweep intensities were used for each experiment, with data from the smallest and largest intensities used for analyses in Figure 2.

*Laser Doppler Vibrometry - CO_2_ sedation experiments*: Mounted mosquitoes were placed inside a rectangular steel chamber (6□×□6□×□2.5Lcm^3^) positioned opposite the laser Doppler vibrometer. The chamber was held in place by a micromanipulator. The chamber side facing the vibrometer contained a glass window, thus enabling the recording of flagellar vibrations from mosquitoes.

Free fluctuation recordings were taken prior to CO_2_ exposure to test the baseline hearing status. CO_2_ was then allowed to flow into the chamber for one minute through the porous membrane floor. CO_2_ was maintained at a constant flow rate of 3Ll/min throughout via a flow regulator (Flowbuddy, Flystuff). Looped free fluctuation recordings were taken whilst CO_2_ was allowed to flow into the chamber to monitor changes to the mosquito’s active hearing system. Once the active system was no longer visible (judged via reference to previous reports)^9^, CO_2_ flow was immediately stopped and a free fluctuation recording was taken. Each mosquito was given 5Lminutes to recover from sedation, after which a final free fluctuation recording was taken.

Mosquito recovery from sedation was judged based on the final free fluctuation recording. Mosquitoes were deemed to have recovered if the best frequency and velocity amplitudes extracted from free fluctuation fits (described below) were altered from the baseline state by less than 20%. Those which did not fit the recovery criteria were excluded from all analyses. These recovery criteria were adopted for all LDV experiments in terms of comparing baseline and final free fluctuations.

*Laser Doppler Vibrometry - Free fluctuation analysis*: As reported previously^9^, a forced damped harmonic oscillator function was fitted to flagellar velocity values obtained by applying Fast Fourier transforms (FFTs) to flagellar velocity amplitudes. The FFTs covered frequencies between 1LHz and 10LkHz, though the function was only fit to values between 101 and 1000Hz due to significant levels of noise below 100Hz.

This function enabled estimation of the flagellar best frequency, as well as other parameters of interest. Analyses were aggregated across individual mosquitoes within a group in order to calculate population estimates and facilitate comparisons across groups.

Sample sizes for each group in the active state were: *dsxF*^+/+^ *XX* = 20; *dsxF*^-/-^ *XX* = 25; *dsxF*^+/+^ *XY* (non-SSO/SSO) = 11/ 21.

Sample sizes for each group in the passive state were: *dsxF*^+/+^ *XX* = 10; *dsxF*^-/-^ *XX* = 15; *dsxF*^+/+^ *XY* (non-SSO/SSO) = 10/ 11.

*Laser Doppler Vibrometry - Apparent mass estimation*: Apparent flagellar mass values are crucial for electrostatic stimulation analyses but can vary significantly across different mosquito sexes and species. We therefore calculated apparent mass values for each group of mosquitoes tested; samples sizes for each group were equal to the total number of passive state mosquito recordings per group.

We fitted the damped harmonic oscillator model described above to velocity spectra obtained from sedated mosquitoes and then calculated the apparent antennal mass based on the formula:

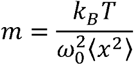

where *m* represents the apparent antennal mass, *k_B_*is the Boltzmann constant (1.38□×□10^−23^□J/K), T the absolute temperature (estimated at approximately 293LK), ω*_0_* the natural frequency of the system and *<x^2^>* the flagellar receiver’s total fluctuation power. ω*_0_* was calculated from the results of the damped harmonic oscillator function fit whilst 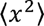 was calculated from:

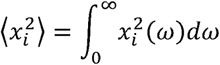

*Laser Doppler Vibrometry - Power gain*: Power gain values were calculated by computing the ratio of total auditory system fluctuation power in active and passive states:

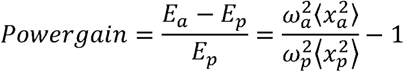

where E_a_ and E_p_ represent the energy contents of the active and passive hearing systems, ω_a_ and ω_p_ the natural frequencies of the active and passive systems and 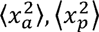 the total fluctuation powers of the active and passive systems respectively. Natural frequencies were calculated based on the results of the damped harmonic oscillator function fits in both active and passive states. Sample sizes for each group were equal to the number of passive state recordings taken per group.

*Laser Doppler Vibrometry - Force step stimulation analysis*: Force step analysis followed previously published protocols by utilising a two-state model of a single transducer population^9^. Displacement data between ±2000nm were analysed for females in order to focus only on the most sensitive transducers.

Sample sizes for each group were: *dsxF*^+/+^ *XX* = 8; *dsxF*^-/-^ *XX* = 8.

*Laser Doppler Vibrometry - Sweeps analysis*: DC removes (with time constants of 0.015) were applied to flagellar displacement and nerve data in Spike2. Averages of flagellar displacement and nerve responses were then created in Spike2 for each stimulus type, with an average of each stimulus also created. These averages were then exported to Matlab for further analysis.

Maximal flagellar displacements were identified using a modified version of the ‘findpeaks’ function in MATLAB. As the change in stimulus frequency was linear, by identifying the timepoint at which this maximum was reached we were able to calculate the corresponding stimulus frequency. A similar procedure was used to identify the maximal nerve response and then estimate the stimulus frequency at which this occurred.

Mechanical sensitivities were calculated for each stimulus intensity by calculating the ratio of maximal flagellar displacement to stimulus magnitude (which remained constant throughout stimulus presentation). A three-parameter sigmoidal function was then fitted to the mechanical sensitivity values at each stimulus intensity, with all fits having calculated R^2^ values ≥0.9. Displacement gains were then calculated by computing the ratio of the maximal and minimal values obtained from the fit.

Sample sizes for each group: *dsxF*^+/+^ *XX* = 8; *dsxF*^-/-^ *XX* = 8; *dsxF*^+/+^ *XY* (SSO) = 10.

*RNA sequencing - Sample collection*: 2-3 day old *dsxF*^+/+^ and *dsxF*^-/-^ *XX* and *XY* adult mosquitoes were placed into glass vials (as described above) in sex- and genotype-sorted groups of five. These vials were then transferred to the incubators described previously. Mosquitoes were entrained using a 12 hour:12 hour light/ dark cycle for three days, with the first hour of light involving a constant increase in light intensity to a maximum and the last hour of light involving a constant decrease in light intensity to a minimum.

On the third day, vials containing mosquitoes were removed from the incubator during the period when light intensity began decreasing. Mosquitoes were immediately transferred to Eppendorfs and frozen in liquid nitrogen. Samples were stored in a -80°C freezer prior to dissections.

During dissections, both the mouthparts and flagellae were first removed from the head. Mosquito pedicels were then also removed from the head using a pair of forceps and transferred to an eppendorff containing 297 μl lysis buffer (InvitrogenTM PureLinkTM RNA Mini Kit) and 3 μl 2-beta-mercaptoethanol. Samples were then submitted for RNAseq total transcriptome Illumina sequencing at the Polo d’Innovazione di Genomica, Genetica e Biologia (PoloGGB).

Sample sizes for each group (number of pedicels): *dsxF*^+/+^ *XX* = 63/76/72; *dsxF*^-/-^ *XX* = 70/72/72; *dsxF*^+/+^ *XY* = 71/64/64.

*RNA sequencing - Data analysis:* Raw read (fastq) files were first subjected to quality control using FastQC and MultiQC, and subsequently classified and quantified against the *An. gambiae* transcriptome (Anopheles-gambiae-PEST_TRANSCRIPTS_AgamP4.12.fa.gz obtained from VectorBase) using Kallisto. Furthermore, STAR was used in performing alignment of reads onto the An. gambiae genome (genome file: Anopheles-gambiae-PEST_CHROMOSOMES_AgamP4.fa.gz; annotation file: Anopheles-gambiae-PEST_BASEFEATURES_AgamP4.12.gtf.gz) to allow for visual inspection of the results using IGV. Quality control and differential expression analysis of read counts were conducted with the R-package DESeq2. The following comparisons were performed:

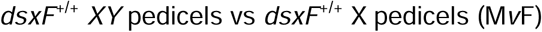

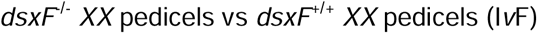

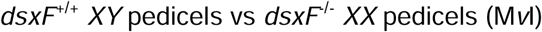

Transcripts were deemed to be significantly differentially expressed if FDR < 0.05 (adjusted p-values from Wald tests). Where information was available, non-annotated transcripts were translated, aligned against the protein database UniProtKB reference proteomes plus Swiss-Prot of Uniprot, and assigned annotation based on homology. Scoring matrix used: BLOSUM-62. Homology thresholds: > 33% query coverage, > 33% identity between alignments, and E-value < 0.0001.

GProfiler^43^ was used to conduct functional enrichment analysis on the genesets obtained by taking the intersections of the results of pairs of comparisons, namely the pairs: 1) MvF and IvF, and 2) MvF and MvI. The analysis was performed with emphasis on upregulated genes. Here upregulated refers to higher expression in the group on the left side of the comparison and characterized by a positive log2(FoldChange) (LFC). For example, if gene X has a positive LFC for the comparison MvF, that gene is upregulated in the male pedicel relative to the female pedicel. A gene ontology was deemed significantly enriched if the FDR < 0.05 (g:SCS threshold). Testes comparisons were also conducted using GProfiler with the same analysis paradigm.

*Statistical analysis*: 20 step and free fluctuation analyses were conducted using Sigmaplot (Systat Software, Inc.). Remaining analyses were completed in Matlab and R. RNAseq analysis used R for differential gene expression, and GProfiler for functional enrichment.

Sample sizes for all experiments were determined via reference to published investigations^9,42^. Within-group variation estimates were calculated when appropriate as part of standard statistical testing.

Statistical tests for normality (Shapiro–Wilk Normality tests with a significance level of p□<□0.05) were first applied to each dataset. Throughout the analyses, all statistical tests used a significance level of p□<0.05.

Flagellar measurements were found to be not normally distributed; ANOVA on ranks tests were thus used for comparisons across the genotypes and sexes.

Data obtained from the free fluctuation analyses were not found to be normally distributed. ANOVA on ranks tests were therefore used for comparisons between genotypes and sexes, with comparisons for males split into SO and quiescent states. Mann-Whitney tests were used for comparisons within a mosquito group between active and passive states.

Data from the single transducer population fits were not found to be normally distributed. ANOVA on ranks tests were therefore used to make comparisons between genotypes.

Best frequencies of both the mechanical and nerve responses to sweeps were found to be normally distributed; t-tests were thus used for comparisons across genotypes and sexes. Paired t-tests were used to compare between the largest and smallest intensity sweeps in terms of both mechanical and nerve best frequencies within a group.

## References

1. Loh, Y.M., Su, M.P., Ellis, D.A., and Andrés, M. (2023). The auditory efferent system in mosquitoes. Front Cell Dev Biol 11, 1123738. 10.3389/fcell.2023.1123738.

2. Andrés, M., Seifert, M., Spalthoff, C., Warren, B., Weiss, L., Giraldo, D., Winkler, M., Pauls, S., and Göpfert, M.C. (2016). Auditory Efferent System Modulates Mosquito Hearing. Curr Biol 26, 2028–2036. 10.1016/J.CUB.2016.05.077.

3. Lapshin, D.N., and Vorontsov, D.D. (2021). Frequency tuning of swarming male mosquitoes (*Aedes communis*, Culicidae) and its neural mechanisms. J Insect Physiol 132. 10.1016/J.JINSPHYS.2021.104233.

4. Loh, Y.M.M., Xu, Y.Y.J., Lee, T.T., Ohashi, T.S., Zhang, Y.D., Eberl, D.F., Su, M.P., and Kamikouchi, A. (2024). Differences in male *Aedes aegypti* and *Aedes albopictus* hearing systems facilitate recognition of conspecific female flight tones. iScience 27, 110264. 10.1016/j.isci.2024.110264.

5. Andrés, M., Su, M.P., Albert, J., and Cator, L.J. (2020). Buzzkill: targeting the mosquito auditory system. Curr Opin Insect Sci 40, 11–17. 10.1016/j.cois.2020.04.003.

6. Diabate, A., and Tripet, F. (2015). Targeting male mosquito mating behaviour for malaria control. Parasit Vectors 8. 10.1186/S13071-015-0961-8.

7. Sawadogo, S.P., Niang, A., Bilgo, E., Millogo, A., Maïga, H., Dabire, R.K., Tripet, F., and Diabaté, A. (2017). Targeting male mosquito swarms to control malaria vector density. PLoS One 12. 10.1371/journal.pone.0173273.

8. Lapshin, D.N. (2013). The auditory system of blood-sucking mosquito females (Diptera, Culicidae): Acoustic perception during flight simulation. Entomol Rev 93, 135–149. 10.1134/S0013873813020012.

9. Su, M.P., Andrés, M., Boyd-Gibbins, N., Somers, J., and Albert, J.T. (2018). Sex and species specific hearing mechanisms in mosquito flagellar ears. Nat Commun. 10.1038/s41467-018-06388-7.

10. Ohashi, T.S., Xu, Y.Y.J., Shigaki, S., Nakamura, Y., Lee, T.-T., Loh, Y.M., Mishiro-Sato, E., Eberl, D.F., Su, M.P., and Kamikouchi, A. (2024). Sexually dimorphic auditory representation in *Aedes aegypti* brains. bioRxiv, 2024.07.07.602439. 10.1101/2024.07.07.602439.

11. Göpfert, M.C., and Robert, D. (2001). Active auditory mechanics in mosquitoes. Proc R Soc Lond B Biol Sci 268, 333–339. 10.1098/RSPB.2000.1376.

12. Georgiades, M., Alampounti, A., Somers, J., Su, M.P., Ellis, D.A., Bagi, J., Terrazas-Duque, D., Tytheridge, S., Ntabaliba, W., Moore, S., et al. (2023). Hearing of malaria mosquitoes is modulated by a beta-adrenergic-like octopamine receptor which serves as insecticide target. Nat Commun 14. 10.1038/s41467-023-40029-y.

13. Hall, A.B., Basu, S., Jiang, X., Qi, Y., Timoshevskiy, V.A., Biedler, J.K., Sharakhova, M. V., Elahi, R., Anderson, M.A.E., Chen, X.G., et al. (2015). A male-determining factor in the mosquito *Aedes aegypti*. Science (1979) 348, 1268–1270. 10.1126/SCIENCE.AAA2850.

14. Krzywinska, E., Dennison, N.J., Lycett, G.J., and Krzywinski, J. (2016). A maleness gene in the malaria mosquito *Anopheles gambiae*. Science 353, 67–69. 10.1126/SCIENCE.AAF5605.

15. Kyrou, K., Hammond, A.M., Galizi, R., Kranjc, N., Burt, A., Beaghton, A.K., Nolan, T., and Crisanti, A. (2018). A CRISPR–Cas9 gene drive targeting doublesex causes complete population suppression in caged *Anopheles gambiae* mosquitoes. Nature Biotechnology 2018 36:11 *36*, 1062–1066. 10.1038/nbt.4245.

16. Camara, N., Whitworth, C., Dove, A., and Van Doren, M. (2019). *Doublesex* controls specification and maintenance of the gonad stem cell niches in Drosophila. Development. 10.1242/dev.170001.

17. Li, M., Kandul, N.P., Sun, R., Yang, T., Benetta, E.D., Brogan, D.J., Antoshechkin, I., C, H.M.S., Zhan, Y., Debeaubien, N.A., et al. (2024). Targeting sex determination to suppress mosquito populations. Elife 12. 10.7554/ELIFE.90199.

18. Degner, E.C., Ahmed-Braimah, Y.H., Borziak, K., Wolfner, M.F., Harrington, L.C., and Dorus, S. (2019). Proteins, Transcripts, and Genetic Architecture of Seminal Fluid and Sperm in the Mosquito *Aedes aegypti*. Molecular & Cellular Proteomics 18, S6–S22. 10.1074/MCP.RA118.001067.

19. Chen, J., Luo, J., Wang, Y., Gurav, A.S., Li, M., Akbari, O.S., and Montell, C. (2021). Suppression of female fertility in *Aedes aegypti* with a CRISPR-targeted male-sterile mutation. Proc Natl Acad Sci U S A 118, e2105075118. 10.1073/PNAS.2105075118.

20. Su, M.P., Georgiades, M., Bagi, J., Kyrou, K., Crisanti, A., and Albert, J.T. (2020). Assessing the acoustic behaviour of *Anopheles gambiae* (s.l.) dsxF mutants: Implications for vector control. Parasit Vectors 13, 1–9. 10.1186/S13071-020-04382-X/TABLES/3.

21. Hammond, A., Galizi, R., Kyrou, K., Simoni, A., Siniscalchi, C., Katsanos, D., Gribble, M., Baker, D., Marois, E., Russell, S., et al. (2016). A CRISPR-Cas9 gene drive system targeting female reproduction in the malaria mosquito vector *Anopheles gambiae*. Nature Biotechnology 2015 34:1 *34*, 78–83. 10.1038/nbt.3439.

22. Wang, Y., Thakur, D., Duge, E., Murphy, C., Girling, I., DeBeaubien, N.A., Chen, J., Nguyen, B.H., Gurav, A.S., and Montell, C. (2024). Deafness due to loss of a TRPV channel eliminates mating behavior in *Aedes aegypti* males. Proc Natl Acad Sci U S A 121. 10.1073/PNAS.2404324121.

23. Boo, K.S., and Richards, A.G. (1975). Fine structure of scolopidia in Johnston’s organ of female *Aedes aegypti* compared with that of the male. J Insect Physiol 21, 1129–1139. 10.1016/0022-1910(75)90126-2.

24. Boo, K.S., and Richards, A.G. (1975). Fine structure of the scolopidia in the johnston’s organ of male *Aedes aegypti* (L.) (Diptera: Culicidae). Int J Insect Morphol Embryol 4, 549–566. 10.1016/0020-7322(75)90031-8.

25. Somers, J., Georgiades, M., Su, M.P., Bagi, J., Andrés, M., Alampounti, A., Mills, G., Ntabaliba, W., Moore, S.J., Spaccapelo, R., et al. (2022). Hitting the right note at the right time: Circadian control of audibility in *Anopheles* mosquito mating swarms is mediated by flight tones. Sci Adv 8, 4844. 10.1126/SCIADV.ABL4844.

26. Cassone, B.J., Kay, R.G.G., Daugherty, M.P., and White, B.J. (2017). Comparative transcriptomics of malaria mosquito testes: Function, evolution, and linkage. G3: Genes, Genomes, Genetics 7, 1127–1136. 10.1534/G3.117.040089/-/DC1.

27. Göpfert, M.C., Humphris, A.D.L., Albert, J.T., Robert, D., and Hendrich, O. (2005). Power gain exhibited by motile mechanosensory neurons in *Drosophila* ears. Proc Natl Acad Sci U S A 102, 325–330. 10.1073/PNAS.0405741102/ASSET/F9BCAE9D-5F4F-45A4-8D5E-541DE79371E8/ASSETS/GRAPHIC/ZPQ0020569170003.JPEG.

28. Boyd-Gibbins, N., Tardieu, C.H., Blunskyte, M., Kirkwood, N., Somers, J., and Albert, J.T. (2021). Turnover and activity-dependent transcriptional control of NompC in the *Drosophila* ear. iScience 24. 10.1016/J.ISCI.2021.102486.

29. Vannini, L., and Willis, J.H. (2016). Immunolocalization of cuticular proteins in Johnston’s organ and the corneal lens of *Anopheles gambiae*. Arthropod Struct Dev 45, 519. 10.1016/J.ASD.2016.10.006.

30. Zhou, Y., Badgett, M.J., Orlando, R., and Willis, J.H. (2019). Proteomics reveals localization of cuticular proteins in *Anopheles gambiae*. HHS Public Access. Insect Biochem Mol Biol 104, 91–105. 10.1016/j.ibmb.2018.09.011.

31. He, D.Z.Z., Lovas, S., Ai, Y., Li, Y., and Beisel, K.W. (2014). Prestin at year 14: progress and prospect. Hear Res 311, 25–35. 10.1016/J.HEARES.2013.12.002.

32. Kavlie, R.G., Fritz, J.L., Nies, F., Göpfert, M.C., Oliver, D., Albert, J.T., and Eberl, D.F. (2015). Prestin is an anion transporter dispensable for mechanical feedback amplification in *Drosophila* hearing. J Comp Physiol A Neuroethol Sens Neural Behav Physiol 201, 51–60. 10.1007/S00359-014-0960-9.

33. Kamikouchi, A., Albert, J.T., and Göpfert, M.C. (2010). Mechanical feedback amplification in *Drosophila* hearing is independent of synaptic transmission. Eur J Neurosci 31, 697–703. 10.1111/J.1460-9568.2010.07099.X.

34. zur Lage, P., Newton, F.G., and Jarman, A.P. (2019). Survey of the Ciliary Motility Machinery of *Drosophila* Sperm and Ciliated Mechanosensory Neurons Reveals Unexpected Cell-Type Specific Variations: A Model for Motile Ciliopathies. Front Genet 10. 10.3389/FGENE.2019.00024.

35. Freeman, E.A., Ellis, D.A., Bagi, J., Tytheridge, S., and Andrés, M. (2024). Perspectives on the manipulation of mosquito hearing. Curr Opin Insect Sci 66, 101271. 10.1016/j.cois.2024.101271.

36. Kalita, A.I., Marois, E., Kozielska, M., Weissing, F.J., Jaouen, E., Möckel, M.M., Rühle, F., Butter, F., Basilicata, M.F., and Keller Valsecchi, C.I. (2023). The sex-specific factor SOA controls dosage compensation in *Anopheles* mosquitoes. Nature 2023 623:7985 *623*, 175–182. 10.1038/s41586-023-06641-0.

37. Krzywinska, E., Ferretti, L., Li, J., Li, J.C., Chen, C.H., and Krzywinski, J. (2021). *femaleless* Controls Sex Determination and Dosage Compensation Pathways in Females of *Anopheles* Mosquitoes. Curr Biol 31, 1084–1091.e4. 10.1016/J.CUB.2020.12.014.

38. Li, X., Jin, B., Dong, Y., Chen, X., Tu, Z., and Gu, J. (2019). Two of the three Transformer-2 genes are required for ovarian development in *Aedes albopictus*. Insect Biochem Mol Biol 109, 92–105. 10.1016/J.IBMB.2019.03.008.

39. Kalita, A.I., Marois, E., Kozielska, M., Weissing, F.J., Jaouen, E., Möckel, M.M., Rühle, F., Butter, F., Basilicata, M.F., and Keller Valsecchi, C.I. (2023). The sex-specific factor SOA controls dosage compensation in *Anopheles* mosquitoes. Nature 623, 175–182. 10.1038/s41586-023-06641-0.

40. Ranson, H., N’Guessan, R., Lines, J., Moiroux, N., Nkuni, Z., and Corbel, V. (2011). Pyrethroid resistance in African anopheline mosquitoes: what are the implications for malaria control? Trends Parasitol 27, 91–98. 10.1016/J.PT.2010.08.004.

41. Johnson, B.J., and Ritchie, S.A. (2016). The Siren’s Song: Exploitation of Female Flight Tones to Passively Capture Male *Aedes aegypti* (Diptera: Culicidae). J Med Entomol 53, 245–248. 10.1093/JME/TJV165.

42. Xu, Y.Y.J., Loh, Y.M.M., Lee, T.T., Ohashi, T.S., Su, M.P., and Kamikouchi, A. (2022). Serotonin modulation in the male *Aedes aegypti* ear influences hearing. Front Physiol 13, 931567. 10.3389/FPHYS.2022.931567.

43. Kolberg, L., Raudvere, U., Kuzmin, I., Adler, P., Vilo, J., and Peterson, H. (2023). g:Profiler—interoperable web service for functional enrichment analysis and gene identifier mapping (2023 update). Nucleic Acids Res 51, W207–W212. 10.1093/NAR/GKAD347.

