## Supplementary figures and images for "Using a female-specific isoform of *doublesex* to explore male-specific hearing in mosquitoes"

### Supplemental Figure 1

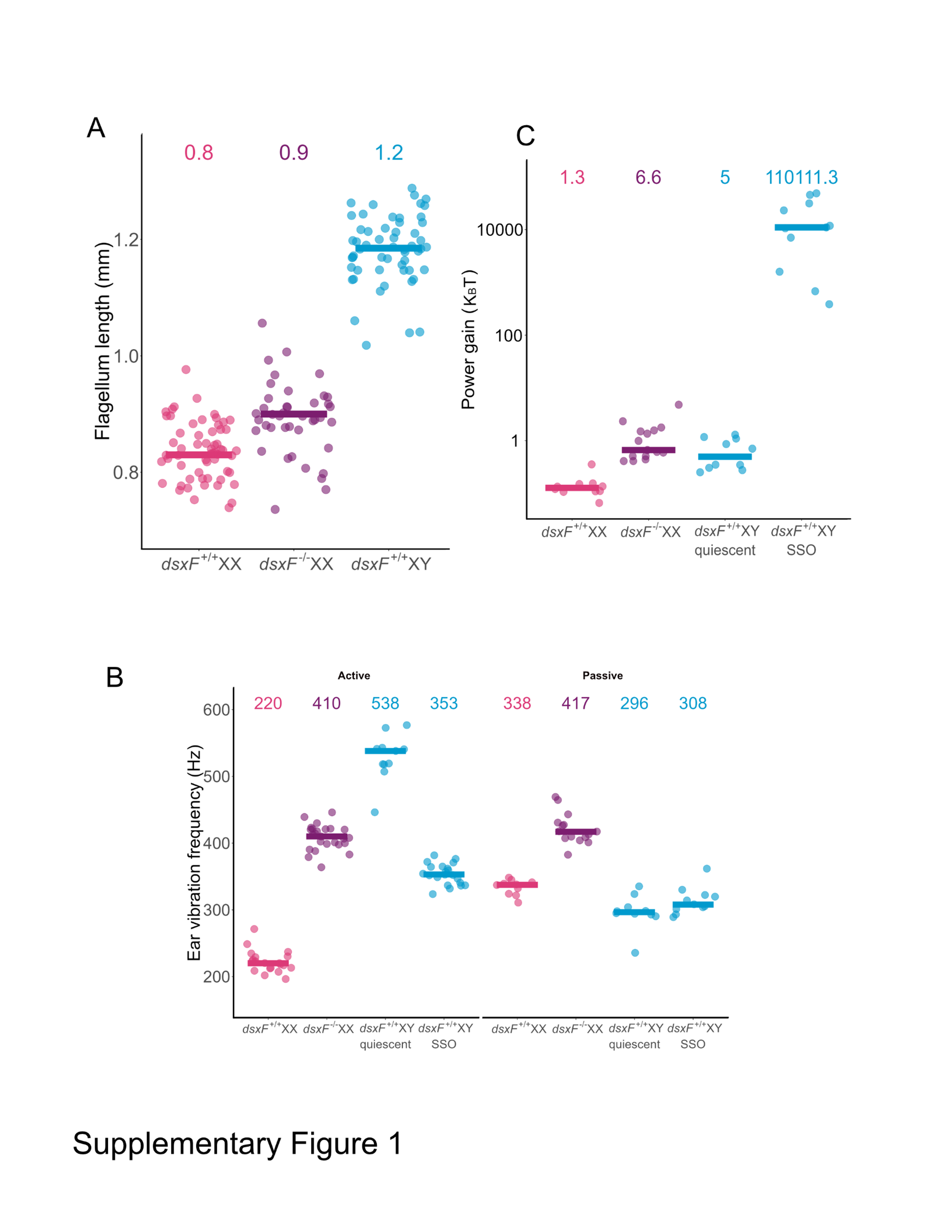

### Supplemental Figure 2

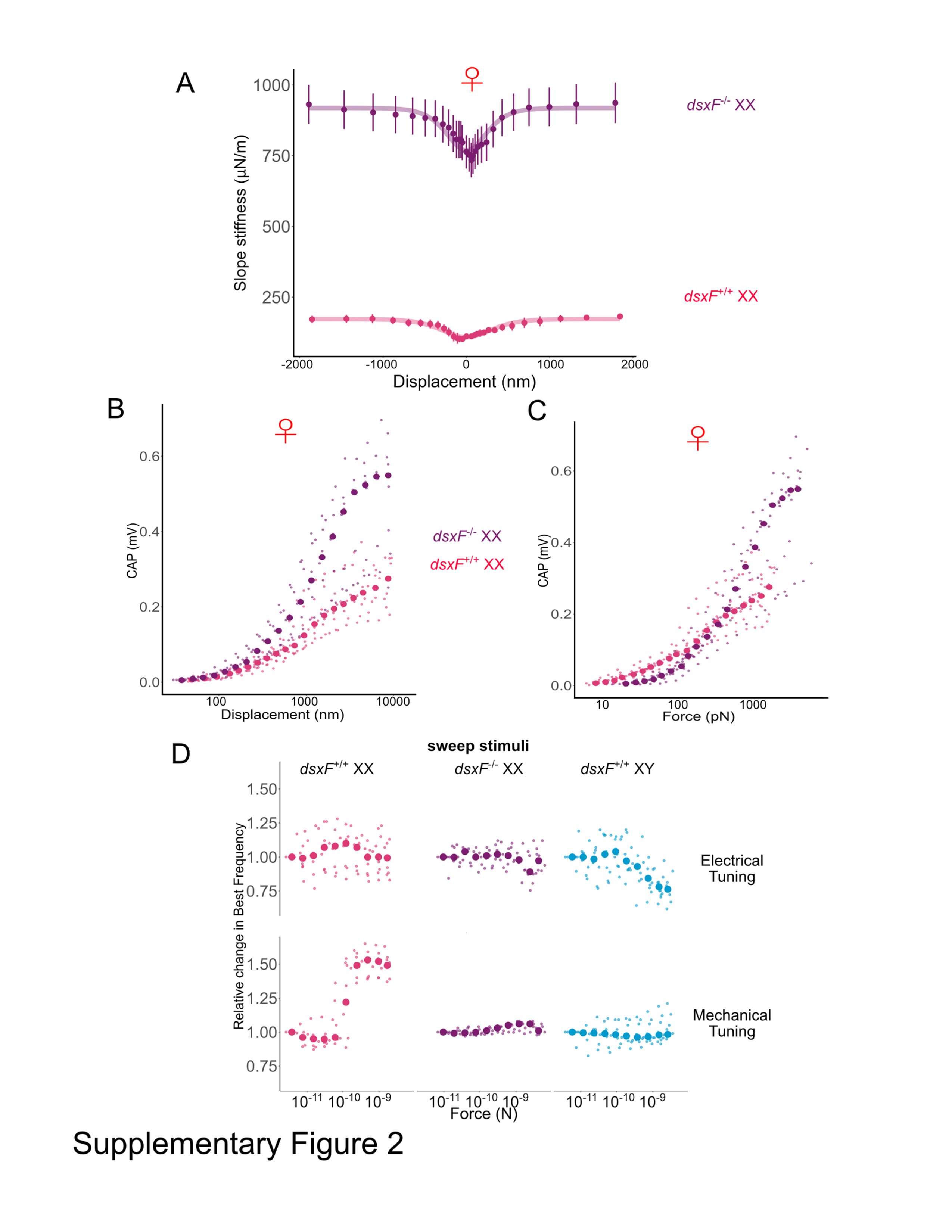

### Supplemental Figure 3

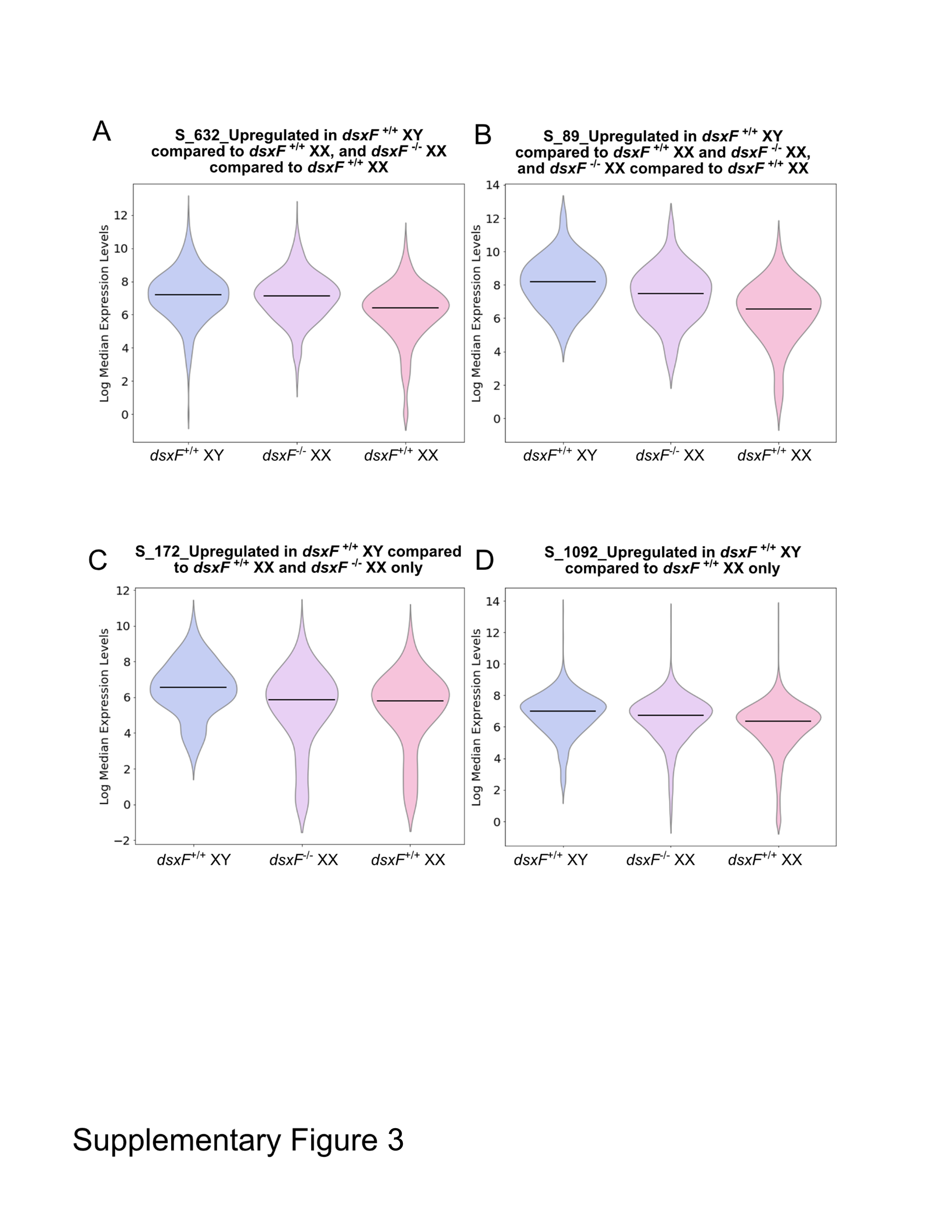

### Supplemental Figure 4

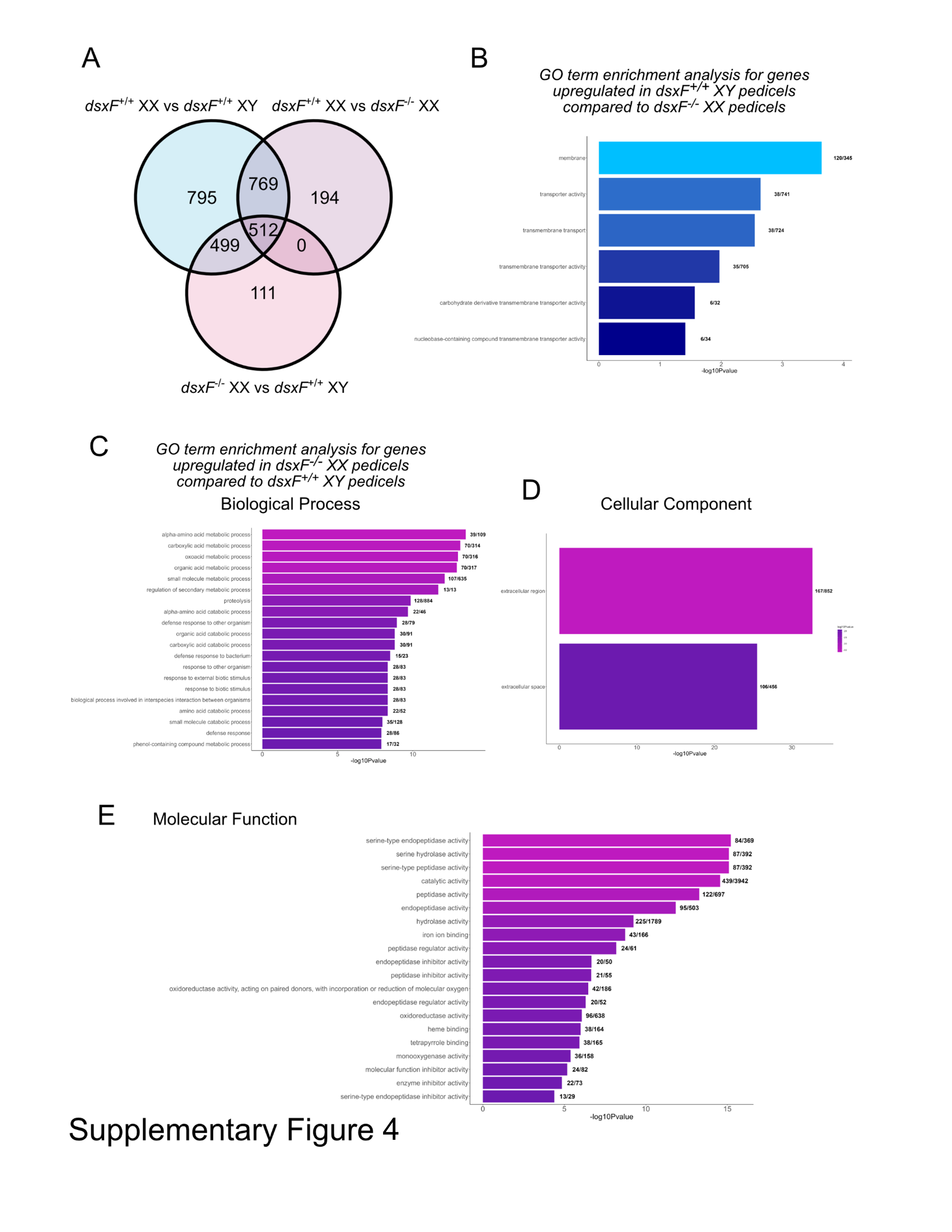
